# Dysfunctions of the paraventricular hypothalamic nucleus induce hypersomnia in human and mice

**DOI:** 10.1101/2021.05.11.443607

**Authors:** Chang-Rui Chen, Yu-Heng Zhong, Shan Jiang, Wei Xu, Zan Wang, Lei Xiao, Wei-Min Qu, Zhi-Li Huang

**Author notes:** These authors contributed equally: Chang-Rui Chen, Yu-Heng Zhong, Shan Jiang and Wei Xu. Correspondence and requests for materials should be addressed to W.-M.Q., C.-R.C. or to Z.-L.H.

## Abstract

Hypersomnolence disorder (HD) is characterized by excessive sleep, which is a common sequela following stroke, infections or tumorigenesis. HD was traditionally thought to be associated with lesions of wake-promoting nuclei. However, lesion of a single, even two or more wake-promoting nucleuses simultaneously did not exert serious HD. The specific nucleus and neural circuitry for HD remain unknown. Here, we observed that three patients with lesions around the paraventricular nucleus of the hypothalamus (PVH) showed hypersomnolence lasting more than 20 h per day and their excessive sleep decreased with the recovery of the PVH area. Therefore, we hypothesized that the PVH might play an essential role in the occurrence of HD. Using multichannel electrophysiological recording and fiber photometry, we found that PVH^vglut2^ neurons were preferentially active during wakefulness. Chemogenetic activation of PVH^vglut2^ neurons potently induced 9-h wakefulness, and PVH^CRH^, PVH^PDYN^ and PVH^OT^ neuronal activation also exerted wakefulness. Most importantly, ablation of PVH^vglut2^ neurons drastically induced hypersomnia-like behaviors (30.6% reduction in wakefulness). These results indicate that dysfunctions of the PVH is crucial for physiological arousal and pathogenesis underlying HD.

## Introduction

Hypersomnolence disorder (HD) is characterized by an irresistible need for sleep and an inability to stay awake during major waking episodes, which results in reduced function and overall worse quality of life and even induces mental diseases, highlighting its public health importance^1^. However, few dysfunctional wake-promoting nuclei have been identified to induce hypersomnia. Therefore, further identification of key hypersomnia control nuclei and neural circuitry represents a common goal for clinicians and researchers.

In the last 100 years, more than 16 wake-promoting nuclei have been identified. Von Economo first proposed a hypersomnia-controlling region located in the posterior hypothalamus from observations of marked somnolence in patients with epidemic encephalitis lethargic [1]. Furthermore, Moruzzi et al. and other studies have revealed that a brainstem ascending reticular activating system (ARAS) is responsible for wakefulness [2–4]. However, cell-body-specific ablation or inhibition of components of the ARAS— including the laterodorsal tegmentum (LDT), basal forebrain (BF), pedunculopontine tegmental nucleus (PPT) cholinergic neurons, dorsal raphe nucleus (DRN) serotonergic neurons, and locus coeruleus (LC) noradrenergic neurons—yields limited alterations in sleep [5–7]. Additionally, the lateral hypothalamic area (LH), parabrachial complex (PB), tuberomammilary nucleus (TMN), paraventricular nucleus of the thalamus (PVT), ventral tegmental area (VTA), and supramammillary nucleus (SUM) have also been demonstrated to be involved in arousal regulation [5, 8–12]. However, among these wake-promoting nuclei, only LH orexinergic and PB glutamatergic neurons have been shown to be related to hypersomnia. Dysfunction of orexinergic neurons in the LH results in narcolepsy and sleep fragmentation [5, 11, 13, 14]; PB glutamatergic neurons are considered to serve as a hub, as they receive afferent chemosensory information and play a role in triggering hypercapnia-induced arousal in obstructive sleep apnea (OSA), whereas ablation of PB glutamatergic neurons decreases hypercapnia-induced arousal [15–17]. The further amazing research found that ablation of LH orexinergic neurons and lateral parabrachial nucleus (L-PBN) glutamatergic neurons have little effect on sleep under baseline conditions, and deletion of vesicular glutamate transporter 2 (vglut2) from the medial PB (MPB) causes only a modest (approximately 20%) reduction in wakefulness [7, 13]. Clinically, patients with Parkinson’s disease (PD), Alzheimer’s disease (AD), Kleine-Levin Syndrome, and idiopathic hypersomnia (IH), in which LH orexinergic and PB glutamatergic neurons are thought to function normally, still show hypersomnolence [18]. Collectively, these results suggest that the key hypersomnolence control nucleus remains unidentified.

Here, we clinically observed that three patients with lesions around the paraventricular nucleus of the hypothalamus (PVH) showed hypersomnolence lasting over 20 h per day and their total sleep time decreased along with the diminished lesion in the PVH area. Considering that the PVH is a highly conserved brain region in species from zebrafish to humans[19]. We used transgenic mice with cutting-edge techniques to reveal the underlying mechanism of this phenotype in patients.

The PVH is located in the ventral diencephalon adjacent to the third ventricle[19]. More than 90% of the PVH consists of glutamatergic neurons, whereas GABAergic neurons are more scarcely represented [20–22]. PVH^vglut2^ neurons co-express corticotropin-releasing hormone (CRH)[23], oxytocin (OT)[24], and prodynorphin (PDYN) [25]. In the present study, we found that activation of PVH^vglut2^, PVH^CRH^, PVH^OT^ and PVH^PDYN^ neurons induced wakefulness. Conversely, ablation or suppression of PVH^vglut2^ neurons caused hypersomnia-like behaviors. Furthermore, photostimulation of PVH^vglut2^→parabrachial complex/ventral lateral septum circuits immediately drove transitions from NREM to wakefulness. Taken together, our findings indicate that the PVH is essential for physiologic arousal and pathogenesis underlying HD.

## Results

### Three patients with lesions around the PVH showed hypersomnia

We observed three clinical cases with central-nervous-system hypersomnia who showed excessive daytime sleepiness (more than 20 h per day), but without any signs of cataplexy, sleep paralysis, or hypnogogic hallucinations. Case a, case b, and case c were diagnosed with stroke, neuromyelitis optical spectrum disorder (NMOSD), and NMOSD recrudescence (NMOSDR), respectively. Brain magnetic resonance imaging (MRI) revealed a lesion restricted to the hypothalamus in each of these three patients. Polysomnographic (PSG) recordings showed that the total sleep time (TST) of each patient was more than 20 h per day, and that stage-two non-rapid eye movement (NREM) sleep was dominant. The patient with NMOSDR (case c) recovered after immunotherapy, as her TST per day decreased and her hypothalamic lesions nearly disappeared by the time of her follow-up MRI (Figure 1A).

**Figure 1.**
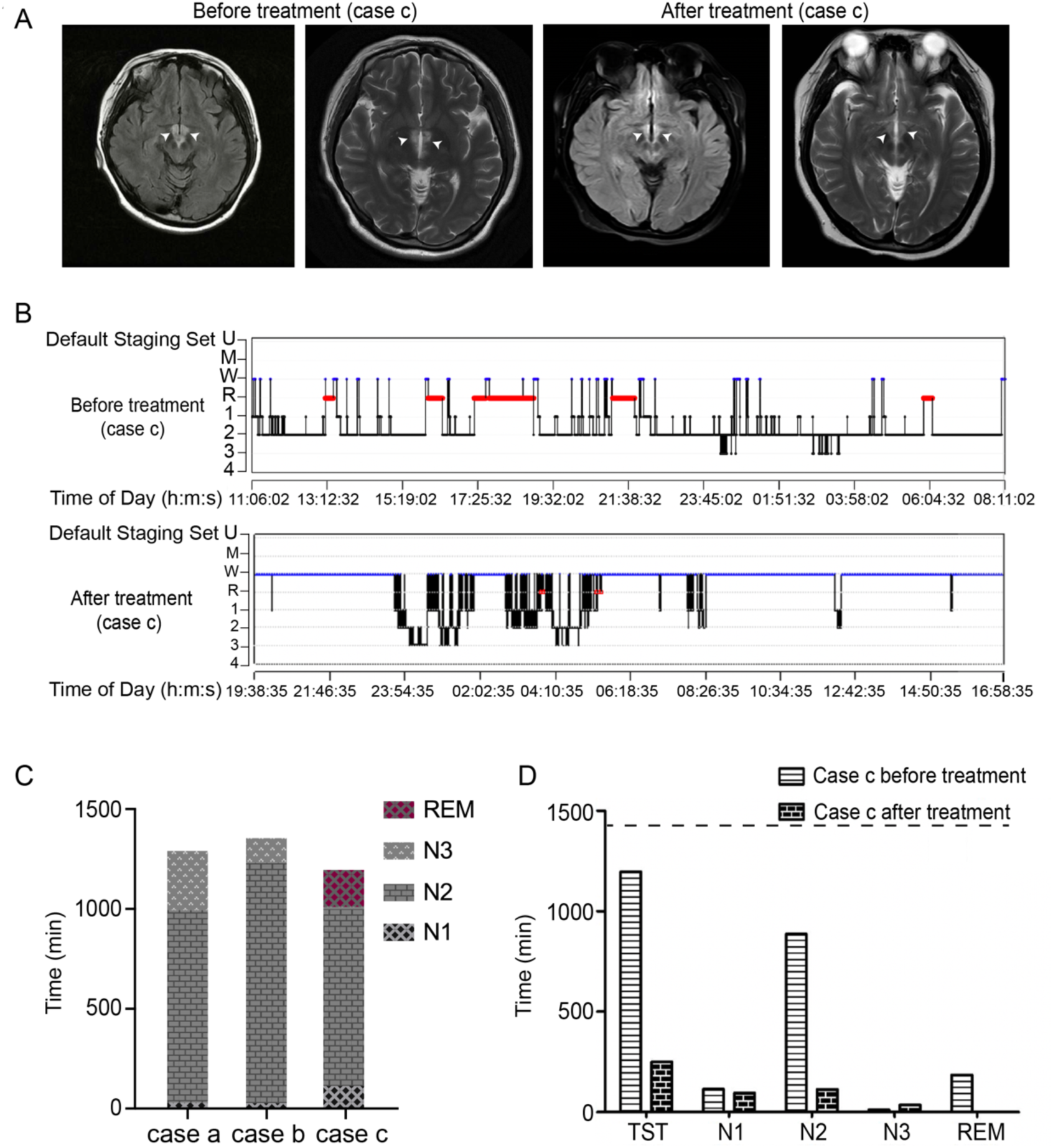
Three cases of patient with lesions in the paraventricular hypothalamic nucleus area showed hypersomnia. **(A)** Fluid-attenuated inversion recovery (FLAIR) showed the lesions of case c was in the bilateral hypothalamus. White arrows indicate the injury sites. **(B)** A sleep-structure chart of case c before treatment and after treatment is shown. Blue lines represent wakefulness, red lines represent REM sleep, and black lines represent NREM sleep (including stages N1, N2, and N3). (**C**) Sleep time of NREM stage 1 (N1), NREM stage 2 (N2), NREM stage 3 (N3) and REM respectively of three cases. The dashed black line indicates 24-h sleep-wake cycle. (**D)** Total sleep time (TST) and the sleep time of NREM stage 1 (N1), NREM stage 2 (N2), NREM stage 3 (N3) and REM respectively of case c before treatment and after treatment during 24h. The dashed black line indicates 24-h sleep-wake cycle.

The clinical information of the included patients is summarized in Table supplement 1. Case a, case b, and case c with hypersomnolence displayed diffusion-weighted imaging (DWI) with a hypersignal in the right hypothalamus (Figure supplement 1A), fluid-attenuated inversion recovery (FLAIR) sequences with a slight hypersignal in the left hypothalamus (Figure supplement 1B), and FLAIR sequences with a slight hypersignal in the bilateral hypothalamus (Figure 1A), respectively. Sleep structure charts showed that the sleep time during stage-two NREM was dominant, accounting for 74.9% of TST in case a, 89.0% in case b, and 74.1% in case c (Figure 1B and 1C and Figure supplement 1C).

After 14 days of antiplatelet aggregation treatment, case a slept less than before, and the muscle strength of her right limb was increased. As for case b and case c, after administration of high-dose (1,000 mg) intravenous methylprednisolone (IVMP) on three consecutive days and oral low-dose methylprednisolone (60 mg) to taper off steroids after IVMP-attack therapy, both patients recovered in terms of decreased sleep time and improved health conditions. Comparing imaging manifestations and sleep quality before and after treatments, we found that the lesioned area in FLAIR sequences became smaller and that TST was decreased (case c, Figures 1A and 1D), suggesting that dysfunctions of the PVH area might act as a central node for the occurrence of HD.

### PVH receives direct inputs from the PVT and PB

Considering that the homologous area of the primate posterior hypothalamus in rodents is around the PVH area, which contains mainly glutamatergic neurons, we examined the role of PVH^vglut2^ neurons in the regulation of wakefulness. We used Cre-dependent rabies virus–mediated monosynaptic retrograde tracing in Vglut2-Cre mice (Figures 2A and 2B), rabies virus infection was clearly detectable at the infection site in Vglut2-Cre mice compared with wild-type littermates, indicating no leakage of viral infection (Figures 2C and 2D). We found that PVH^vglut2^ neurons received direct inputs from the paraventricular thalamic nucleus (PVT), parabrachial nucleus (PB), zona incerta (ZI) and ventrolateral periaqueductal gray (VLPAG) (Figure 2E and Figure supplement 2), which were reported involving in sleep-wake control [9, 15–17, 26, 27], suggesting that the PVH might act as a key central node for sleep-wake regulation.

**Figure 2.**
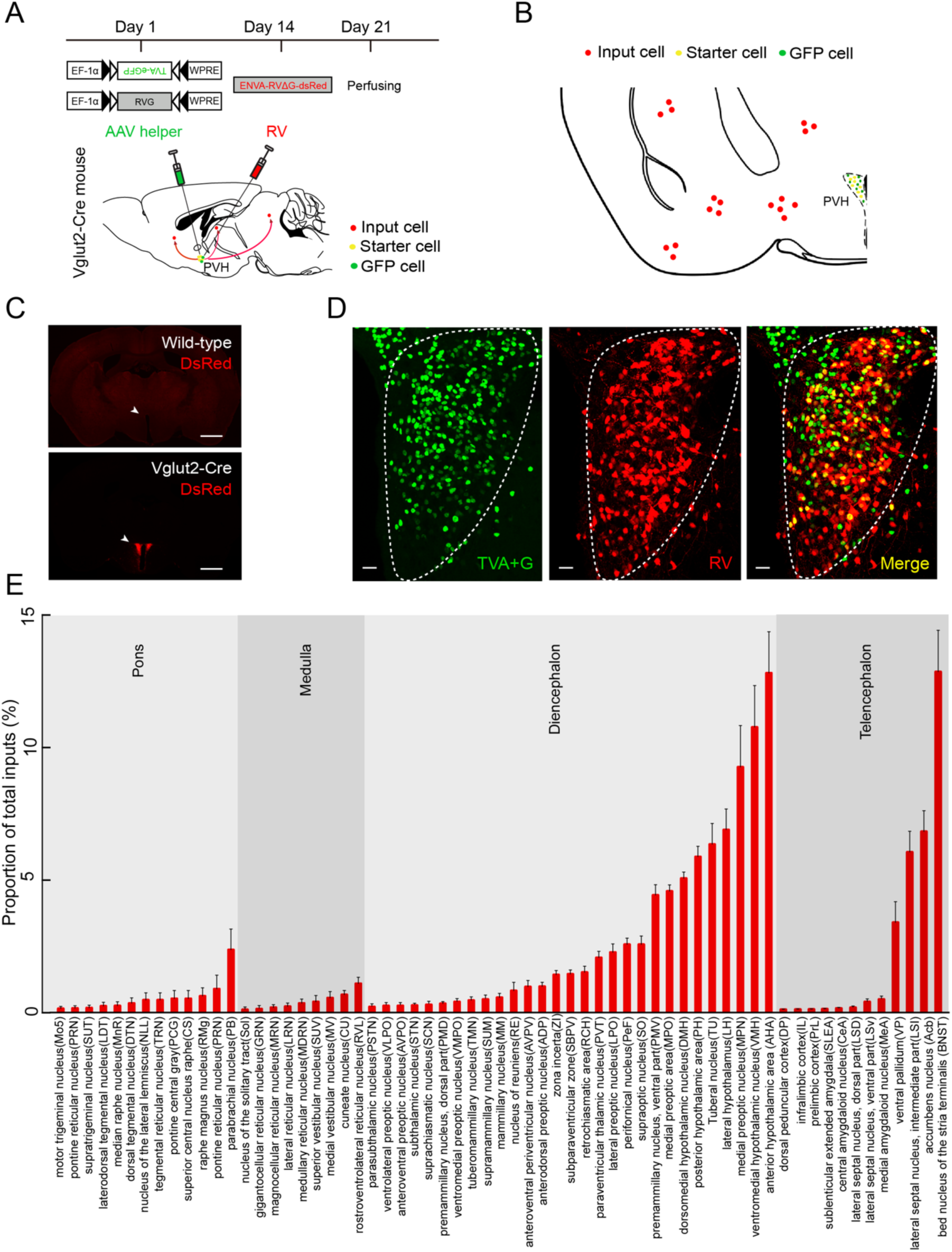
RV retrograde tracing in PVH^vglut2^ neurons. **(A)** Experimental procedure of RV retrograde tracing in PVH^vglut2^ neurons. Top: experimental timeline for injecting Cre-dependent helper viruses into the PVH of Vglut2-Cre mice. Bottom: a schematic of microinjection into the PVH in Vglut2-Cre mice. **(B)** A schematic drawing showed a brain section of the anatomical site and viral infection in the PVH. **(C)** Typical fluorescence images showing RV only labelled in Vglut2-Cre mice (top) rather than in wild-type mice (bottom). Scale bar: 500 μm. White arrows indicated the PVH area. **(D)** Fluorescence images showing that the starter cells (yellow) labelled by both helper viruses (green) and RV (red) were restrictedly infected in the unilateral PVH in a Vglut2-Cre mouse. Scale bar: 50 μm. **(E)** Percentage of whole-brain, monosynaptic inputs to PVH^vglut2^ neurons. Total number of mice counted here was four. Red column with different length represented the proportion (or intensity) of inputs.

### PVH^vglut2^ neurons are preferentially active during wakefulness and necessary for the control of natural wakefulness

We next performed *in-vivo* multichannel electrophysiological recordings to monitor the spike firing of individual PVH neurons in freely behaving mice (Figure 3A). PVH neurons exhibited a higher firing rate during wakefulness than during sleep (Figures 3B and 3C). Neuronal firing rate in the PVH gradually decreased before sleep onset and increased during transitions from sleep to wakefulness (Figure 3D–3F). At the onset of behavioral arousal from NREM sleep, the mean firing rate reached 13.52 Hz (Figure 3D). We next performed *in-vivo* fiber photometry to investigate the real-time activity of PVH^vglut2^ neurons across spontaneous sleep–wake cycles in freely moving mice. The recording mode for fiber photometry and the expression of the Cre-dependent AAVs expressing the fluorescent calcium indicator, GCaMP6f (AAV-EF1α-DIO-GCaMP6f), in the PVH of Vglut2-Cre mice are shown in figure supplements 3A and 3B. PVH^vglut2^ neuronal activities during wakefulness were significantly higher than those during NREM sleep (Figure supplement 3C–3E). Collectively, these electrophysiological results clearly indicate a mechanistic framework for the activity-dependent participation of PVH neurons in the regulation of sleep and wakefulness.

**Figure 3.**
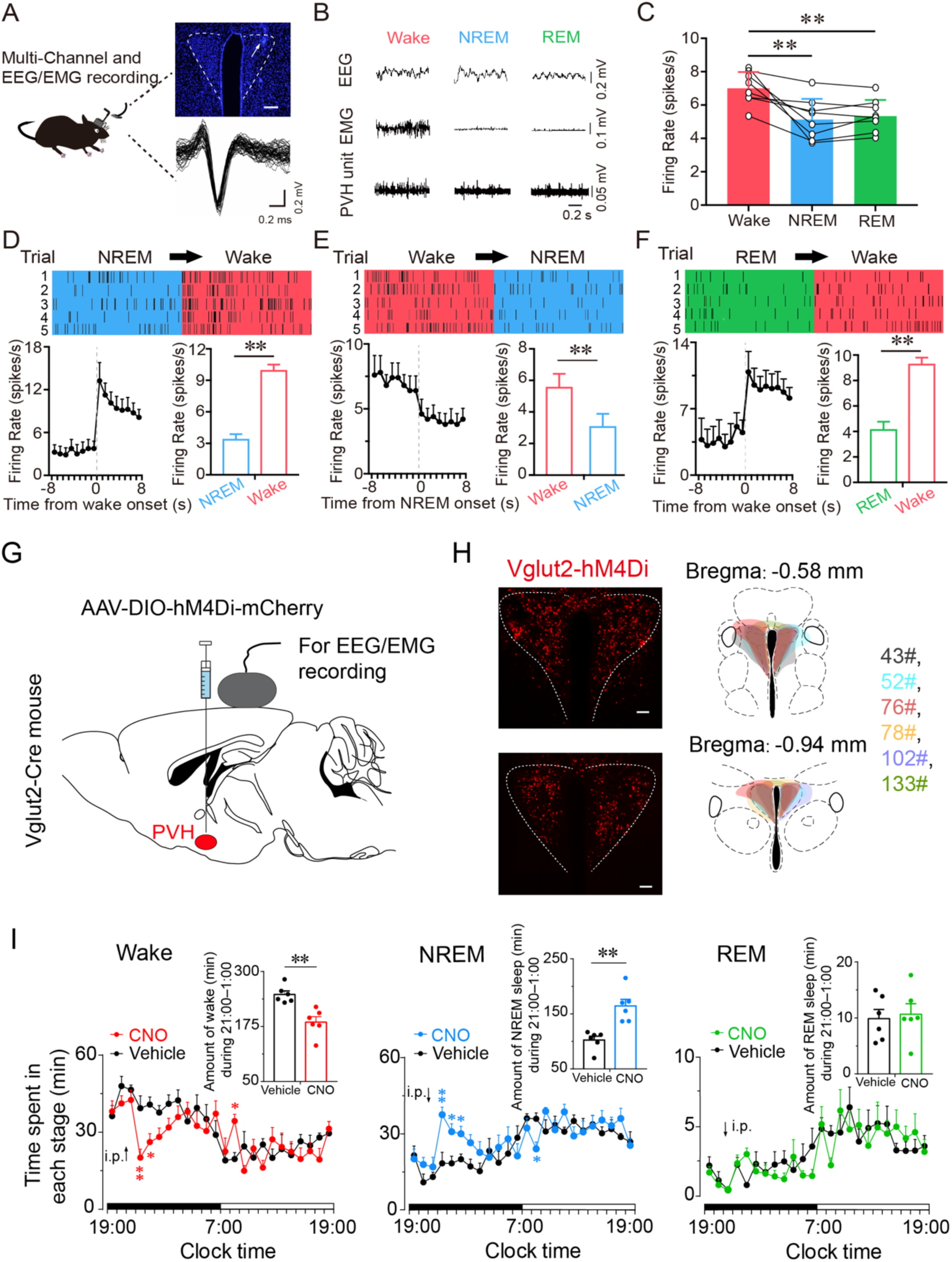
PVH^vglut2^ neurons are preferentially active during wakefulness and inhibition of PVH^vglut2^ neurons decreased NREM sleep. **(A)** Schematic conFigureuration of *in-vivo* multichannel electrophysiological recordings. Upper panel: A brain slice from a mouse with electrodes implanted in the PVH. White arrow indicates the electrode track. Scale bar: 200 μm. Lower panel: Waveforms from a recorded PVH neuron. **(B)** EEG/EMG and PVH multi-unit recording traces during wakefulness, NREM sleep, and REM sleep. **(C)** Average firing rates of PVH neurons during each state (n=3 mice, one-way RM ANOVA followed by LSD post hoc tests; F_2,14_ =12.51, *P* [NREM vs wake] < 0.001, *P* [wake vs NREM] < 0.01, *P* [NREM vs REM] = 0.613). **(D-F)** Firing rates of PVH neurons during state transitions: wake-to-NREM (**D**), NREM-to-wake (**E**), and REM-to-wake transitions (**F**). Top: Example rastergrams of a PVH neuron during five trials of different state transitions. Bottom left: Average firing rate during the state-transition period. Bottom right: Average firing rate during 8 s before and after state transitions (*P* [wake–NREM] < 0.01, *P* [wake–NREM] < 0.01, *P* [REM–wake] < 0.01, paired t test). **(G)** Expression of AAV injection site in the PVH of Vglut2-Cre mice. **(H)** Left panel: location of hM4Di expression in the PVH^vglut2^ neurons. Right panel: drawings of overlay mCherry expressing sites in the PVH of Vglut2-Cre mice (n = 6, indicated with different colors). Scale bars: 200 μm. **(I)** Time-course changes in NREM sleep, wakefulness, and REM sleep after administration of vehicle or CNO in mice expressing hM4Di in PVH^vglut2^ neurons (n = 6, repeated-measures ANOVA; F_1,10_= 21.95 [wake], 7.68 [NREM], 29.23 [REM]). Inset: Total time spent in each stage after vehicle or CNO injection (n = 6, paired t test). Data represented the mean ± SEM (\**P* < 0.05, \*\**P* < 0.01).

To determine whether PVH^vglut2^ neurons are necessary for natural wakefulness, we inhibited PVH^vglut2^ neurons with AAV constructs encoding engineered Gi-coupled hM4D receptor (AAV-EF1α-DIO-hM4D(Gi)-mCherry, Figures 3G and 3H). Chemogenetic inhibition of PVH^vglut2^ neurons induced 3 h increase of NREM sleep compared with that of vehicle. At the beginning of the dark phase (ZT15; 21:00), CNO injection resulted in a 64.0% increase in NREM sleep during the 5 h post-injection period, which was accompanied by a 26.0% decrease in wakefulness (Figure 3I). These results suggested that the PVH^vglut2^ neurons might act as a critical role in regulating sleep.

### Activation of PVH^vglut2^ neurons significantly increases wakefulness

Next, we investigated the activation effect of PVH^vglut2^ neurons in freely moving mice on wakefulness regulation by injecting adeno-associated virus (AAV)-EF1α-double-floxed inverse-orientation (DIO)-hM3D(Gq)-mCherry into the PVH, respectively (Figures 4A and 4B). At the beginning of the light phase (zeitgeber time 3 [ZT3]; 9:00), chemogenetic activation of PVH^vglut2^ neurons caused a potent increase in wakefulness lasting approximately 9 h and concomitantly decreased both NREM and REM sleep (Figure 4C). CNO administration (3 mg/kg) resulted in a 139.97% increase in total wakefulness, as well as 81.63% and 94.51% reduction in NREM and REM sleep, respectively, during the 9-h post-injection period (Figure 4D). Compared with vehicle injection, chemogenetic activation of PVH^vglut2^ neurons significantly increased electroencephalographic (EEG) low delta power (0.25–1.00 Hz) and decreased high delta power (1.25–4.75 Hz) (Figure 4F). No sleep rebound followed the long-lasting wakefulness, as indicated by no change in the time spent in NREM sleep during the following dark period (19:00–07:00; Figure 4E). Besides, there is no significant difference in the EEG power density of NREM sleep during the day (7:00–18:00) before/after the day of CNO injection (Figure 4G). Similarly, CNO injection during the dark period also significantly increased wakefulness (Figure supplement 4), further demonstrating that activation of PVH^vglut2^ neurons prolonged arousal even during the dark (active) period.

**Figure 4.**
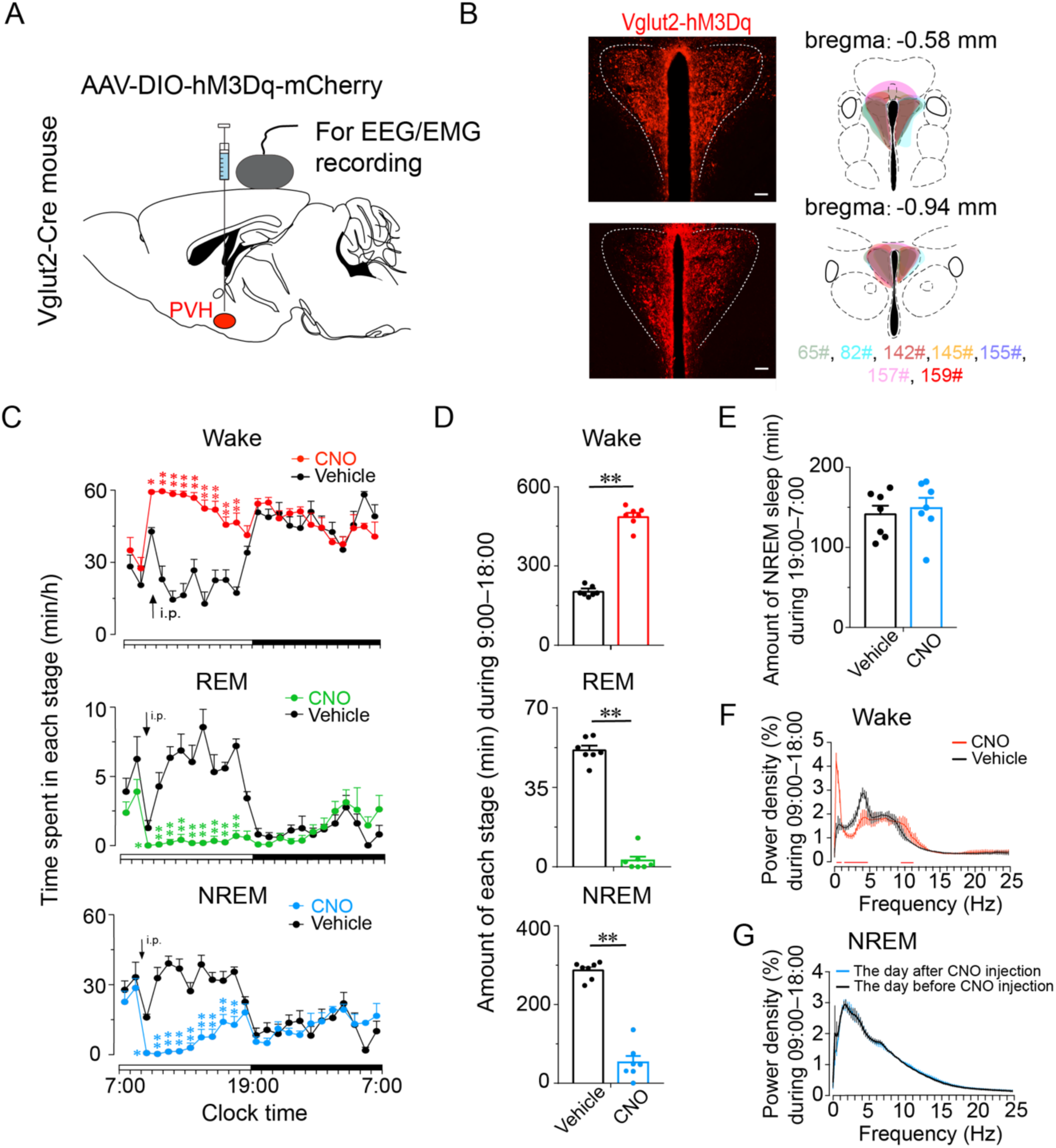
Chemogenetic activation of PVH^vglut2^ increases wakefulness. **(A, B)** Expression of AAV injection site in the PVH of Vglut2-cre mice. Drawings of overlay mCherry expressing sites in the PVH of Vglut2-cre mice (n = 6, indicated with different colors). **(C)** Time-course changes in wakefulness, NREM sleep, and REM sleep after administration of vehicle or CNO in mice expressing hM3Dq in PVH^vglut2^ neurons (n = 7, repeated-measures ANOVA; F_1,12_ = 87.09 [wake], 63.61 [NREM], 612.30 [REM]; *P* < [wake], *P* < 0.001 [NREM], *P* < 0.001 [REM]). **(D)** Total time spent in each stage after vehicle or CNO injection to Vglut2-IRES-cre mice (n = 7, paired t test; *P* < 0.001 [wake], *P* < 0.001 [NREM], *P* < 0.001 [REM]). **(E)** Total time spent in NREM sleep during the dark period after vehicle or CNO injection (n = 7, *P* > 0.05, paired t test). **(F)** EEG power density of wakefulness during 9 h after vehicle or CNO injection (n = 5; *P* < 0.05, paired t test). **(G)** EEG power density of NREM sleep during the day (7:00–18:00) before/after the day of CNO injection (n = 5, *P* > 0.05, paired t test). Data represent the mean ± SEM (\**P* < 0.05, \*\**P* < 0.01, n.s. means no significant difference).

Considering the potent wake-promoting effect of PVH^vglut2^ neurons and the millisecond-scale control of neuronal activity through optogenetic manipulation, we next employed optogenetic methods to elucidate the causal role of the PVHVglut2 neurons in controlling wakefulness. We stereotaxically injected AAVs expressing channelrhodopsin-2 (AAV-DIO-ChR2-mCherry) into the PVH (Figure supplement 5A). Functional expression of ChR2 was verified by in-vitro electrophysiology (Figure supplement 5B). Next, we applied optical blue-light stimulation (10 ms, 20 Hz, 20–30 mW/mm2) after the onset of stable NREM or REM sleep during the light phase (Figure supplement 5C). Optical stimulation of PVH^vglut2^ neurons during NREM sleep reliably induced transitions to wakefulness in a frequency-dependent manner (Figure supplement 5D). Analysis of the probability of transitions between each pair of sleep-wake states showed that optical stimulation significantly enhanced the probability of wakefulness, along with a complementary decrease in the probability of NREM or REM sleep (Figure supplement 5E). To test whether these neurons also contributed to the maintenance of wakefulness, photostimulation was given for 1 h during the light period (09:00–10:00). Sustained activation of PVH^vglut2^ neurons via semi-chronic optical stimulation (10-ms blue-light pulses at 20 Hz for 25 s, every 60 s for 1 h) significantly increased the amount of wakefulness in ChR2-mCherry mice compared with that of the baseline control between 09:00 and 10:00 (12.3 ± 1.8 min at baseline vs. 48.6 ± 1.5 min after stimulation, n = 5; Figure supplement 5F). These findings demonstrate that optogenetic activation of PVH^vglut2^ neurons potently enhanced both the initiation and maintenance of wakefulness.

### PVH^vglut2^ neurons exert wakefulness via PVH^OT^, PVH^PDYN^ and PVH^CRH^ neurons

Next, we further explored arousal-promoting roles of subtype neurons of PVH^vglut2^ neurons (PVH^OT^, PVH^PDYN^ and PVH^CRH^ neurons) and found that chemogenetic activation of PVH^OT^ and PVH^PDYN^ neurons both induced increased wakefulness (49.4% and 75.7%, respectively) and decreased NREM sleep (53.9% and 88.5%) that lasted for 1 h (Figures 5C and 5F). Similarly, chemogenetic activation of PVH^CRH^ neurons caused a potent increase in wakefulness lasting approximately 3 h and concomitantly decreased both NREM and REM sleep. CNO administration (3 mg/kg) induced a 75.8% increase in wakefulness and a 67.7%, 46% reduction in NREM and REM sleep during 3-h post-injection period (Figure 5I).

**Figure 5.**
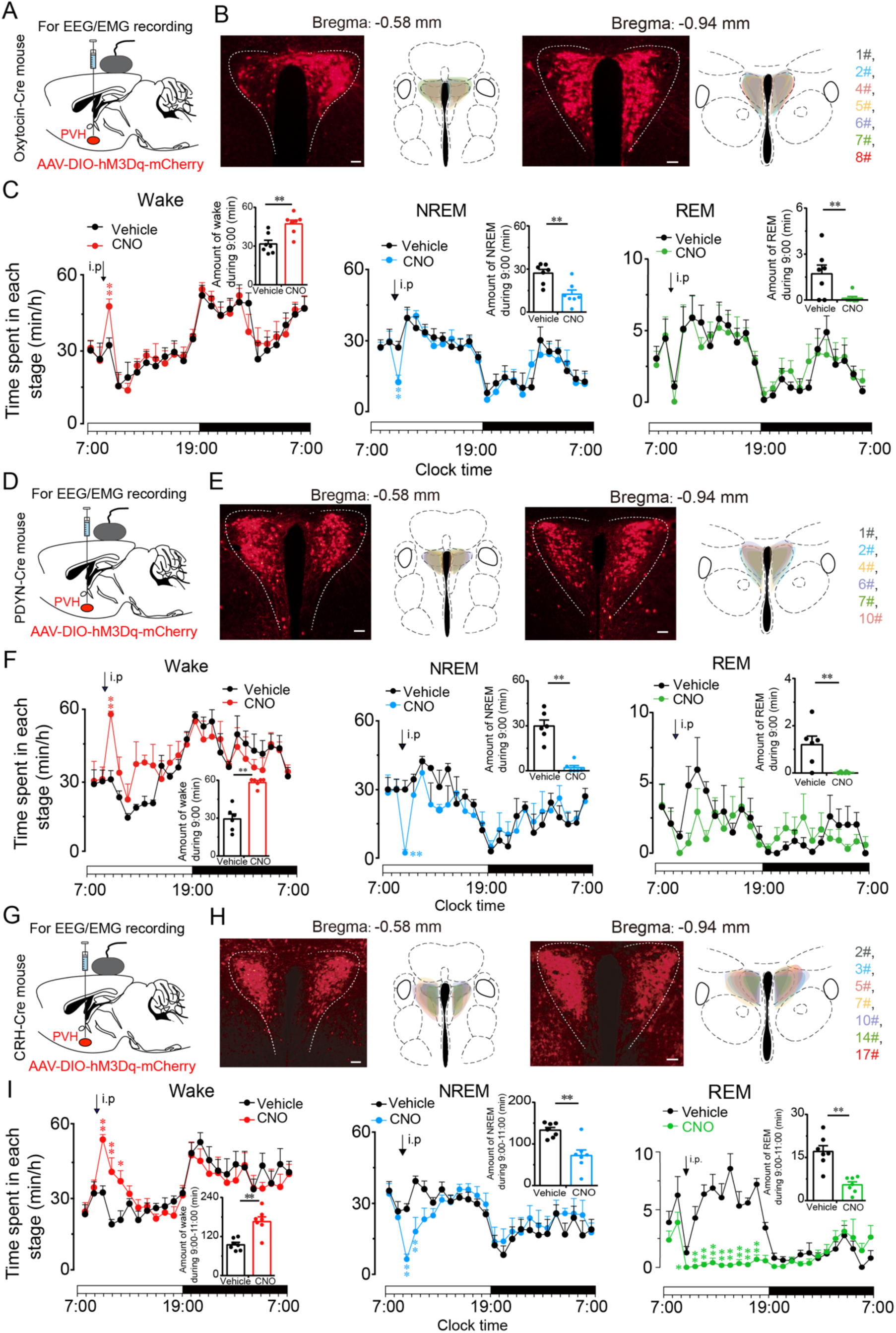
PVH^vglut2^ neurons exert wakefulness via PVH^OT^, PVH^PDYN^ and PVH^CRH^ neurons. **(A, D, J)** Schematic drawing of the chemogenetic experiment in Oxytoxin-Cre mice, PDYN-Cre mice and CRH-Cre mice. (**B, E, H)** Location of hM3Dq expression in PVH^OT^, PVH^PDYN^ and PVH^CRH^ neurons. Right panel: drawings of overlay mCherry expressing sites in the PVH. Scale bars: 200 μm. (**C, F, I)** Time-course changes in wakefulness, NREM sleep, and REM sleep after administration of vehicle or CNO in mice expressing hM3Dq in PVH^OT^ (**c**, n = 7, repeated-measures ANOVA; F_1, 12_= 0.41 [wake], 0.74 [NREM], 0.02 [REM]; *P* < 0.05 [wake], *P* < 0.05 [NREM], *P* < 0.05 [REM]), PVH^PDYN^ neurons (**f**, n = 6, repeated-measures ANOVA; F_1, 10_= 0.28 [wake], 0.38 [NREM], *0.1* [REM]; *P* < 0.05 [wake], *P* < 0.05 [NREM], *P* < 0.05 [REM]) and PVH^CRH^ neurons (**i**, n = 7, repeated-measures ANOVA; F_1, 12_= 0.06 [wake], 0.01 [NREM], 0.83 [REM]; *P* < 0.05 [wake], *P* < 0.05 [NREM], *P* < 0.05 [REM]). Inset: Total time spent in each stage after vehicle or CNO injection to mice expressing hM3Dq in PVH^OT^ neurons (**c**, n = 7, paired t test; *P* = 0.10 [wake], 0.10 [NREM], 0.39 [REM]), PVH^PDYN^ neurons (**f**, n = 6, paired t test, *P* < 0.01 [wake], *P* < 0.01 [NREM], *P* < 0.01 [REM]) and PVH^CRH^ neurons (**i**, n = 5, paired t test; *P* = 0.003 (wake), *P* = 0.002 (NREM), *P* = 0.1 (REM)). \*\**P* < 0.01.

### PVH^vglut2^ neurons promote wakefulness via PB and LSv connections

We next sought to determine the downstream targets by which PVH^vglut2^ neurons promote wakefulness. Specifically, AAV-hSyn-DIO-eGFP constructs were injected into the PVH of Vglut2-cre mice. We found that PVH^vglut2^ neurons mainly projected to two neuroanatomical sites: the PB and LSv may involving in exerting wakefulness (Figure supplement 6). To identify the neuronal circuits mediating the wake-promoting effect of PVH^vglut2^ neurons, ChR2 was expressed in the PVH with optic fibers targeting terminals in the PB or LSv (Figure supplement 7A and 7E). Optogenetic stimulation (10-ms pulses at 10 Hz for 2 s) of the ChR2-expressing PVH terminals evoked excitatory postsynaptic currents (EPSCs) in most of the patch-recorded PB (n = 6 cells, Figure supplement 7B) or LSv neurons (n = 8 cells, Figure supplement 7F). Moreover, 20-Hz stimulation of the bilateral PB or LSv induced a shorter transition from NREM sleep to wakefulness (latency for PB: 1.0 ± 0.8 s, latency for LSv: 1.2 ± 0.9 s) compared with that in the control (Figure supplement 7C and 7G). Analysis of the probability of transitions between each pair of sleep-wake states showed that optical stimulation significantly enhanced the probability of wakefulness, along with a complementary decrease in the probabilities of NREM and REM sleep (Figure supplement 7D and 7H). These results demonstrate that PVH→PB and PVH→LSv circuits mediated the wakefulness-controlling effect of PVH^vglut2^ neurons.

### Ablation of PVH^vglut2^ neurons induces hypersomnia-like behaviors

To further assess the functional importance of PVH^vglut2^ neurons controlling physiological wakefulness, we specifically ablated these neurons by bilaterally microinjecting AAV-EF1a-DIO-taCasp3-TEVp into the PVH region of Vglut2-Cre mice. This construct expressed a designer pro-caspase-3 (pro-taCasp3) in the PVH, the activation of which causes apoptosis (Figures 6A and 6B). Compared with that of the control group, mice that underwent PVH^vglut2^ neuronal ablation showed a 28.6% decrease in the amount of wakefulness and a 74.7% increase in the amount of NREM sleep during the dark period. Similarly, ablation of PVH^vglut2^ neurons induced a 20.8% reduction in wakefulness and 30.6% increase in NREM sleep across an entire 24-h light/dark cycle (Figures 6C and 6D). These results indicate that PVH^vglut2^ neurons are necessary for wake regulation under physiological conditions, and that dysfunction of these neurons may induce hypersomnia.

**Figure 6.**
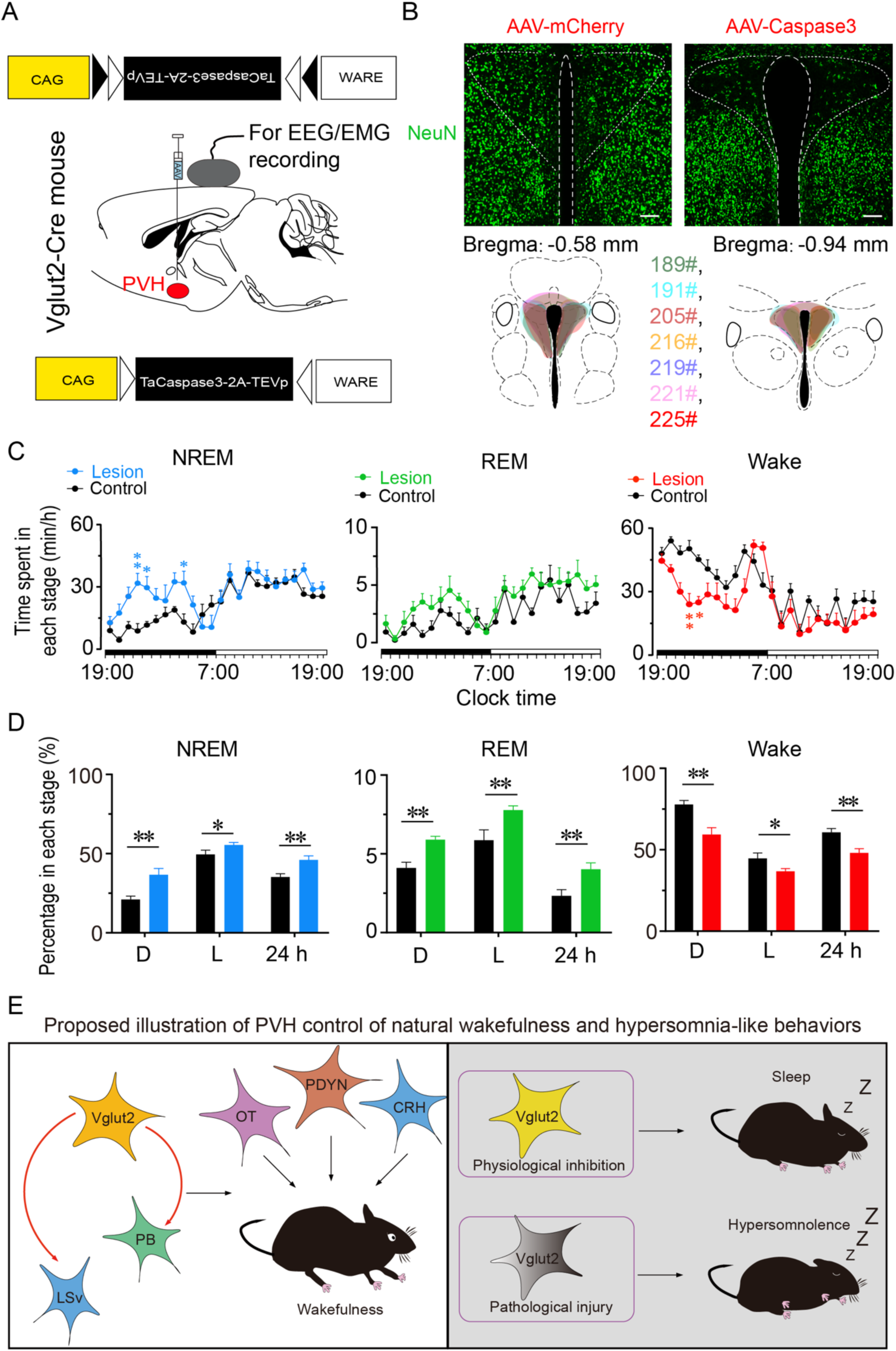
Ablation of PVH^vglut2^ neurons induces hypersomnia-like behaviors. **(A)** Expression of an AAV injection site in the PVH of Vglut2-IRES-cre mice. **(B)** Upper panel: Image showing NeuN (neuron-specific nuclear protein) staining from a control mouse (left) and a mouse with a PVH lesion (right). Lower panel: Drawings of superimposed ablation areas in the PVH of Vglut2-IRES-Cre mice (n = 7, indicated with different colors). Scale bars: 200 μm. **(C)** Time spent in each stage across the 24-h sleep-wake cycle. L, light phase; D, dark phase. (control group, n = 6; lesion group, n = 7, repeated-measures ANOVA; F_1,10_= 2.88 [wake], 2.90 [NREM], 0.06 [REM]; **(D)** Percentages in different sleep-wake stages across the 24-h sleep-wake cycle (unpaired t test; NREM, dark phase, t_11_=3.27, *P* <0.01, light phase, t_11_=2.32, *P* =0.04, 24 h, t_11_=3.27, *P* <0.01; REM, dark phase, t_11_=2.94, *P* =0.01; light phase, t_11_=2.88, *P* =0.05; 24 h, t_11_=4.36, *P* <0.01; Wake, dark phase, t_11_=3.61, *P*<0.01; light phase, t_11_=2.24, *P* =0.05; 24 h, t_11_=3.56, *P* <0.01). **(E)** Model of the PVH control of wakefulness and hypersomnia-like behaviors. Left: Increasing levels of PVH^vglut2^ neurons projecting to the PB and the LSv, thereby activating the PB and the LSv neurons to control wakefulness. Besides, activation of PVH^vglut2^ subtype neurons (PVH^OT^, PVH^PDYN^ and PVH^CRH^ neurons) also induced wakefulness. Top right: decreased activity of PVH^vglut2^ neurons leads to sleep. Bottom right: The impairment of PVH^vglut2^ neurons in neurological diseases may be associated with hypersomnia. Data represented the mean ± SEM (\**P* < 0.05, \*\**P* < 0.01).

## Discussion

Adequate wakefulness is essential for life and survival. In the present study, we identified the PVH as a critical hypothalamic nucleus for the pathogenesis underlying HD in both humans and rodents. In previous study, 15% reduction in baseline wakefulness is considered significant [5, 28]. Lu *et al* have reported that lesion of the PPT and the ventral sublaterodorsal nucleus (vSLD) results in a 20–30% reduction in baseline wakefulness [5]. However, bidirectional chemogenetic manipulations that inhibit the PPT or activate SLD neurons have been shown to have little influence on baseline sleep [29, 30]. In our present study, three patients with lesions mostly around the PVH showed hypersomnolence lasting above 20 h per day. PSG recordings from these patients showed that stage-two NREM was strikingly dominant, indicating that these patients slept stably and were not easily awakened. Importantly, we found that following recovery from injury around the PVH in one of these patients, the proportion of stage-two NREM sleep decreased, and this patient was concomitantly better able to stay awake. Furthermore, ablation of PVH^vglut2^ neurons in mice induced a 30.6% reduction in wakefulness across the 24-h light/dark cycle, highlighting the significance of PVH^vglut2^ neurons in maintaining wakefulness and preventing hypersomnia. Besides, in our murine experiments, no sleep rebound was seen after PVH^vglut2^-activation-induced enhancement of wakefulness. This finding is in accordance with previous studies using chemogenetics to specifically activate wake-promoting neuronal populations [8, 30–32] and indicates that chemogenetic activation of wake-promoting neuronal populations does not enhance the homeostatic drive for sleep. Taken together, our present findings provide evidence of the sufficient and necessary wake-promoting action of PVH^vglut2^ neurons in preventing hypersomnia.

The PVH is composed of abundant, diverse, and functionally distinct groups of neuroendocrine neurons, including CRH, OT, and PDYN neurons[23, 24, 33, 34]. The PVH is estimated consist of approximately 56,000 neurons in humans[35], of which 25,000 neurons express OT, 21,000 neurons express PDYN[34, 36, 37] and 2,000 neurons express CRH[38]. Over 90% of PVH^CRH^ neurons express vglut2 mRNA[22]. PVH^CRH^ neurons regulate stress, fear, and immune responses, as well as neuroendocrine and autonomic functions [23, 39–42]. There is mounting evidence that exposure to various stressors induces CRH and OT release into the peripheral circulation[43]. However, OT exhibits some opposing actions to those of CRH. CRH serves as the starting point and main driver of the hypothalamic-pituitary-adrenal (HPA) axis[23, 44, 45], while OT inhibits the activity of the HPA axis[46]. PVH^CRH^ neurons orchestrate stress-related behaviors, such as grooming, fear, rearing, and walking[45, 47], while PVH^OT^ neurons modulate reward circuits and play a role in mitigating stress responses[43, 48]. In the present study, our results showed that PVH^CRH^, PVH^OT^ and PVH^PDYN^ neurons play important roles in the regulation of wakefulness. Considering that PVH^CRH^ play role in the circadian regulation of wakefulness[49] and optogenetic stimulation of PVH^CRH^ neurons can simulate stress-induced insomnia[50], our results provide further evidence that PVH^CRH^ neurons play an important role in stress-related insomnia. PVH^PDYN^ neurons are key regulators of satiety, and PVH^PDYN^ neurons project to the PB[34]. We found that chemogenetic activation of PVH^PDYN^ neurons induced short-duration wakefulness; hence, dysfunction of PVH^PDYN^ neurons may represent one of the causes of sleep-related eating disorders (SREDs).

HD is one of the most common symptoms in many neurological disorders and mental diseases, including PD, AD, IH, OSA, Huntington’s disease (HD), major depressive disorder (MDD), bipolar disorder (BD), Kleine-Levin syndrome, and depression[18, 51]. A robust reduction in the number of PVH neurons has been found in MDD, BD, AD, and PD patients[52–54]. Thus, our present findings provide evidence that dysfunction of the PVH may contribute to the occurrence of hypersomnia in these diseases. Unfortunately, the clinical component of our present study was limited by missing data from case a and case b after their treatments due to these patients declining to undergo post-treatment PSG monitoring; however, these two patients reported that their post-treatment sleep durations had returned to normal levels. Besides, we observed that the reduction of wakefulness time measured via PSG monitoring showed an earlier recovery compared to that revealed via MRI, suggesting that PSG monitoring may be a more direct and sensitive tool for predicting the prognosis.

In conclusion, our results indicate that dysfunctions of the PVH is crucial for the pathogenesis underlying HD and PVH^vglut2^ and their subtype neurons (PVH^OT^, PVH^PDYN^ and PVH^CRH^ neurons) are critical for wakefulness.

## Methods

### Human studies

#### Subjects

Patients with somnolence were included from the Neurology Department at the First Hospital of Jilin University.

#### Assessments

##### Magnetic resonance imaging (MRI)

All patients completed MRI scanning with a 3.0-T MRI scanner (Siemens Trio Tim 3.0T), including axial T1-weighted imaging (T1WI), T2-weighted imaging (T2WI), diffusion-weighted imaging (DWI), fluid-attenuated inversion recovery (FLAIR), and coronal/sagittal scanning, with a slice thickness of 5 mm, and a slice spacing of 1 mm.

##### Polysomnography (PSG)

All patients were monitored for 24 h in the sleep center of our hospital via PSG recordings (Compumedics, Australia). Simultaneous monitoring of the following was performed: EEG, electromyogram (EMG), bilateral anterior tibial EMG. All data were analyzed according to the revised interpretation criteria of sleep stages and related events issued by the American Academy of Sleep Medicine (AASM) version 2.1.

#### Rodents studies

##### Animals

Vglut2-Cre mice were obtained from Jackson Laboratory (Bar Harbor, Maine, USA). CRH-Cre mice were obtained from the Shanghai Model Organisms Center. Mice were housed in a soundproof room at an ambient temperature of 24 ± 0.5°C, with a relative humidity of 60 ± 2%. A 12-h light/dark cycle (100 Lux, light on at 07:00) was automatically controlled[55]. Food and water were available *ad libitum*. Male heterozygous mice at 6–8 weeks of age were used for all experiments. All animal experiments were approved by the Medical Experimental Animal Administrative Committee of Shanghai. All experimental procedures involving animals were approved by the Animal Experiment and Use Committee of Fudan University (20150119 - 067).

#### Preparation of viral vectors

The AAVs of serotype rh10 for AAV-hSyn-DIO-hM3Dq-mCherry, AAV-hSyn-DIO-hM4Di-mCherry, AAV-hSyn-DIO-ChR2-mCherry, AAV-hSyn-DIO-mCherry, and AAV-CAG-FLEX-taCasp3-TEVp were used. AAV vectors were packaged into serotype 2/9 vectors, which consisted of AAV2 ITR genomes coupled with AAV9 serotype capsid proteins. The final viral concentrations of the transgenes were in the range of 1–5 × 10^12^ viral particles/mL.

#### Surgery and injection of viral vectors

All mice were anesthetized with chloral hydrate (360 mg/kg, i.p.) for surgical procedures and were placed in a stereotaxic apparatus (RWD, Shenzhen, China). The skin above the skull was cut, a burr hole was made, and a small craniotomy was performed above the PVH. AAV constructs were slowly injected (30 nL/min) into the bilateral PVH (70 nL for each position; AP = -0.5 mm; ML = ±0.2 mm; DV = -4.2 mm) for PSG recordings and brain-slice electrophysiology, or were unilaterally injected into the PVH for neuronal tracing. The glass pipette was left in the brain for an additional 10 min following injections and was then slowly withdrawn. All mice were implanted with electrodes for EEG and EMG recordings that were used for *in-vivo* tests at four weeks after injections under anesthesia of chloral hydrate (intraperitoneal, 360 mg/kg). The implant consisted of two stainless steel screws (1 mm in diameter), and EEG electrodes were inserted through the skull (+1.5 mm anteroposterior; -2.0 mm mediolateral from bregma or lambda), while two flexible silver wires were inserted into the neck muscles. The electrodes were attached to a mini-connector and were fixed to the skull with dental cement. The scalp wound was sutured, and the mouse was when kept in a warm environment until it resumed normal activity.

#### Polysomnographic recordings and analysis

After a 2–3-week recovery period, each mouse was individually housed in a recording chamber and habituated to the recording cable for 2–3 days before electrophysiological recordings. Simultaneous EEG/EMG recordings were carried out with a slip ring so that movement of the mice would not be restricted. For experiments using designer receptors exclusively activated by designer drugs (DREADDs), the recordings started at 07:00 (i.e., at the beginning of the light period), and each mouse received either vehicle or CNO (3 mg/kg, C2041, LKT) treatment for two consecutive days at 09:00 (inactive period) or 21:00 (active period). As previously described[9, 56], EEG/EMG signals were amplified and filtered (0.5–30 Hz for EEG, 40–200 Hz for EMG), and were then digitized at 128 Hz and recorded with SleepSign software (Kissei Comtec, Nagano, Japan). Sleep–wake states were automatically classified into 4-s epochs as follows: wakefulness was considered to have desynchronized EEG and high levels of EMG activity, NREM sleep was considered to have synchronized, high-amplitude, low-frequency (0.5–4 Hz) EEG signals in the absence of motor activity; and REM sleep was considered to have pronounced theta-like (4–9 Hz) EEG activity and muscle atonia. All scoring was automated based on EEG and EMG waveforms in 4-s epochs for both chemogenetic and optogenetic studies.

#### Optogenetic stimulation

Before the testing day, mice were given one day to adapt to optical fiber cables (0.8-m long, 200-μm diameter; RWD) that were placed inside the implanted fiber cannulae. On the testing day, 473-nm laser pulses (10 ms, 20 Hz) were delivered via an optic cable (Newton Inc., Hangzhou, China) using a pulse generator. Light pulse trains were generated via a stimulator (SEN-7103, Nihon Kohden, Japan) and delivered through an isolator (ss-102J, Nihon Kohden). For acute photostimulation, each stimulation epoch was applied at 20 s after identifying a stable NREM or REM sleep event via real-time online EEG/EMG analysis. Light pulse trains (5-ms pulses of various frequencies and durations) were programmed and conducted during the inactive period. For chronic photostimulation, programmed light pulse trains (5-ms pulses at 20 Hz for 10 s and at 30-s intervals for 1 h) were used. The 473-nm laser stimulation was performed from 09:00 to 10:00. Baseline EEG/EMG recordings were acquired at the same time of day on the previous day prior to laser stimulation. Sleep–wake cycle parameters (e.g., durations of NREM sleep, REM sleep, and wakefulness, as well as sleep–wake transitions) were scored over an entire hour for each mouse. After receiving photostimulation, mice were sacrificed at 30 min after the final stimulation for subsequent c-Fos staining.

#### In-vitro electrophysiological recordings

At 3–4 weeks after AAV-ChR2 injections, Vglut2-Cre mice were anesthetized and transcardially perfused with ice-cold slicing buffer containing the following (in mM): 213 sucrose, 26 NaHCO_3_, 10 glucose, 0.1 CaCl_2,_ 3 MgSO_4_, 2.5 KCl, 1.25 NaH_2_PO_4_, 2 sodium pyruvate, and 0.4 ascorbic acid. The buffer was saturated with 95% O_2_ and 5% CO_2_. Brains were then rapidly removed, and acute coronal slices (300 μm) containing the PVH were cut using a vibratome (Leica VT 1200S, Nussloch, Germany). Next, slices were transferred to a holding chamber containing normal recording artificial cerebrospinal fluid (aCSF) containing the following (in mM): 119 NaCl, 26 NaHCO_3_, 25 glucose, 2.5 KCl, 2CaCl_2_, 1.25 NaH_2_PO_4_, and 1.0 MgSO_4_. After being transferred, splices were allowed to recover for 30 min at 32°C. Then, slices were maintained at room temperature for at least 30 min before recordings. During recordings, slices were transferred to and submerged in a recording chamber in which oxygenated aCSF was continuously perfused.

Expression of ChR2 was confirmed by visualization of mCherry fluorescence in PVH neuronal somata and axonal terminals. Neurons were identified and visualized with an upright microscope (Olympus, Japan) equipped with differential contrast optics, including a 40× water-immersion objective lens (BX51WI, Olympus). Images were detected with an infrared-sensitive CCD camera (OptiMOS, Q-imaging). Patch-clamp recordings were performed with capillary glass pipettes filled with an intrapipette solution containing the following (in mM): 130 potassium gluconate, 10 KCl, 10 Hepes, 0.5 EGTA, 4 ATP-Mg, 0.5 GTP-Na, and 10 phosphocreatine, adjusting to a pH of 7.2–7.4 with KOH.

Whole-cell patch-clamp recordings were obtained using a MultiClamp 700B amplifier (Molecular Device, Union City, CA, USA) and a Digidata 1440A A/D converter (Molecular Device). Signals were sampled at 10 kHz and filtered at 2 kHz. Data were acquired and analyzed using pClamp 10.3 software (pClamp, Molecular Devices). ChR2 stimulation was evoked using 470-nm light. In voltage-clamp experiments, the holding potential was -70 mV. When needed, 20 μM of 2,3-dihydroxy 6-nitro-7-sulfamoyl-benzoquinoxaline-2,3-dione (NBQX, 1044, Tocris Bioscience, UK), 25 μM of d-(-)-2-amino-5-phosphonopentanoic acid (D-AP5, 0106, Tocris Bioscience, UK), and 10 μM of SR95531 (SR, ab144487, Abcam Biochemicals, UK) were added to block N-methyl-D-aspartic acid receptor (NMDA), α-Amino-3-hydroxy-5-methyl-4-isoxazolepropionic acid (AMPA), and gamma aminobutyric acid A (GABAA) receptors, respectively.

#### Fiber photometry

Following AAV-EF1α-DIO-GCaMP6f injections, an optical fiber (125-μm outer diameter, 0.37 numerical aperture; Newdoon, Shanghai) was placed in a ceramic ferrule and was inserted toward the PVH. Fiber photometry[57] uses the same fiber to both excite and record from GCaMP in real time. After surgery, mice were individually housed for at least 10 days to recover. Fluorescent signals were acquired with a laser beam passed through a 488-nm excitation laser (OBIS 488LS; Coherent), reflected off a dichroic mirror (MD498; Thorlabs), focused by an objective lens (Olympus), and coupled through a fiber collimation package (F240FC-A, Thorlabs) into a patch cable connected to the ferrule of an upright optic fiber implanted in the mouse via a ceramic sleeve (125 μm O.D.; Newdoon, Shanghai). GCaMP6 fluorescence was bandpass filtered (MF525–39, Thorlabs) and collected by a photomultiplier tube (R3896, Hamamatsu). An amplifier (C7319, Hamamatsu) was used to convert the photomultiplier-tube current output to voltage signals, which were further filtered through a low-pass filter (40-Hz cut-off; Brownlee 440). The photometry voltage traces were down-sampled using interpolation to match the EEG/EMG sampling rate of 512 Hz via a Power1401 digitizer and Spike2 software (CED, Cambridge, UK).

Photometry data were exported to MATLAB R2018b mat files from Spike2 for further analysis. We segmented the value of the fluorescent change (ΔF/F) by calculating (F – F_0_)/F_0_, where F_0_ is the baseline of the fluorescent signal. We recorded data for 3–5 h per mouse for the analysis of sleep–wake transitions to calculate the averaged calcium signal of ΔF/F during all times of vigilant states. For analyzing state transitions, we determined each sleep-wake transition and calculated ΔF/F in a ±40-s window around that time point.

#### Firing rate analysis

Electrophysiological data were filtered with a band-pass filter (300–6,000 Hz) to obtain neuronal spikes. Single-unit activities were sorted according to a threshold and shape detector using principal component analysis via Offline Sorter software (Plexon Co, USA). The first two principal components of each spike on the two-dimensional plot of detected spike events were extracted. Waveforms with similar principal components were clustered via a K-means sorting method. The isolated cluster was considered as a single unit recorded from the same neuron. Spikes with inter-spike intervals < 2 ms were discarded. Cross-correlation histograms were used to eliminate cross-channel artifacts. NeuroExplorer software (version 5.0) was used for producing firing-rate rastergrams, and Prism (version 7.0) was used for producing firing-rate histograms.

#### Histology and immunohistochemistry

For dual immunostaining with c-Fos and mCherry, mice were deeply anesthetized with chloral hydrate (400 mg/kg) and were perfused with phosphate-buffered saline (PBS) followed by 4% PFA in 0.1-M phosphate buffer. The brain was then dissected and fixed in 4% PFA at 4°C overnight. Fixed samples were sectioned into 30-μm coronal sections using a freezing microtome (CM1950, Leica, Germany). For immunohistochemistry, the floating sections were washed in PBS and were then incubated in the following primary antibodies in PBS containing 0.3% Triton X-100 (PBST) at 4°C: anti-rabbit c-Fos (1:10000 for 48 h); primary antibody (Millipore); and anti-mouse NeuN (1:1000 for 12 h; MAB377, Millipore). Primary antibodies were washed five times with PBS before incubation with secondary antibodies at room temperature for 2 h (Alexa 488, 1:1000; abcam). Finally, the sections were mounted on glass slides, dried, dehydrated, and cover-slipped. Fluorescent images were collected with a confocal microscope (Nikon AIR-MP).

#### Statistical analysis

All data are expressed as the mean ± standard error of the mean (SEM). Sample sizes were chosen based on previous studies [56, 58]. Two-way repeated-measures analysis of variance (ANOVA) was used to perform group comparisons with multiple measurements. Paired and unpaired t tests were used for single-value comparisons. One-way ANOVA was used to compare more than two groups, followed by *post-hoc* Tukey tests for multiple pairwise comparisons. Prism 7.0 (GraphPad Software, San Diego, CA, USA) was used for all statistical analyses. A two-tailed P < 0.05 was considered statistically significant.

## Acknowledgements

This study was supported in part by the National Natural Science Foundation of China (81671317, 31530035, 81420108015, 31671099, 31871072, 31571103, 81701305, 81970727 and 31900738), the National Basic Research Program of China (2015CB856401), Shanghai Pujiang Program (19PJ1401800) and ZJLab, Program for Shanghai Outstanding Academic Leaders (to Zhi-Li Huang). We are grateful to Hui Dong, Ze-Ka Chen, Ya-Nan Zhao, Peng-Fei Xu, Ming-Jie Yang, Fan Yang, and An-Shu Chen for technical assistance.

## Author contributions

Chang-rui Chen, Zhi-Li Huang conceived and designed the experiments. Yu-Heng Zhong and Shan Jiang performed the experiments and analyzed the data. Wei Xu performed patch-clamp electrophysiology. Lei Xiao performed multichannel electrophysiological recordings. Zan Wang collected the clinical data. Chang-Rui Chen, Yu-Heng Zhong, Shan Jiang wrote the manuscript, and all of the authors helped with the revision of the manuscript.

## Supplementary Materials

**Figure supplement 1.**
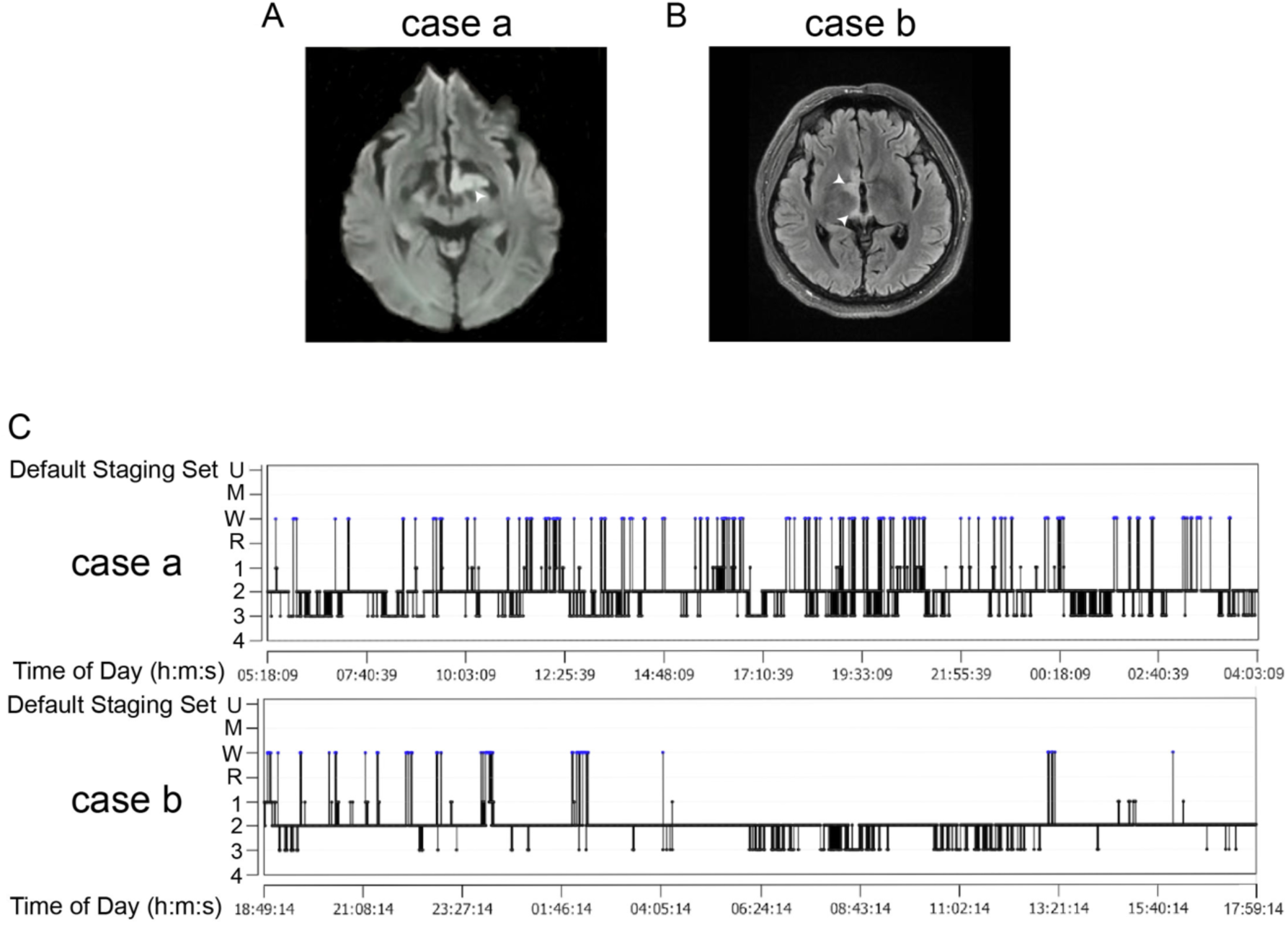
Two cases of patients with injury in the paraventricular hypothalamic nucleus area. **a, b** Diffusion-weighted imaging (DWI) showed strong signals in the left hypothalamus (**A**). The injury was in the right hypothalamus and showed a slight hyper signal in the fluid-attenuated inversion recovery (FLAIR) image (**B**). White arrows indicate the injury sites. **(C)** A sleep-structure chart is shown. The sleep time of stage-two NREM was dominant, accounting for 74.9% of the total sleep time in case a and 74.1% in case b. Blue lines represent wakefulness, red lines represent REM sleep, and black lines represent NREM sleep (including stages N1, N2, and N3).

**Figure supplement 2.**
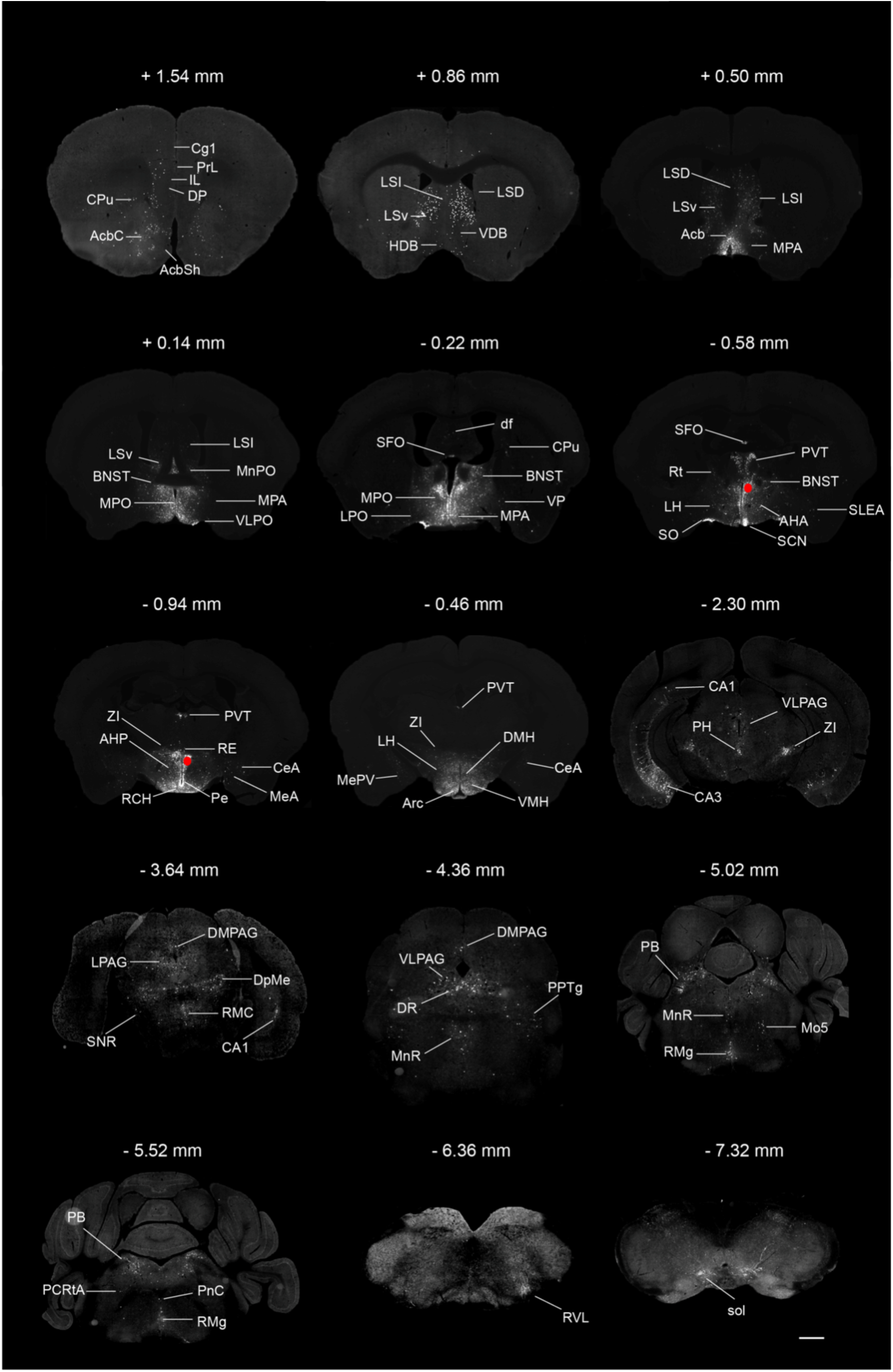
Monosynaptic inputs to PVH^vglut2^ neurons. Typical brain sections showing the source of monosynaptic afferents to PVHvglut2 neurons. The RV infection sites were marked by red rounds. Abbreviations of the brain regions used are the following: Cg1, cingulate Cg1, cortex, area 1; PrL, prelimbic cortex; IL, infralimbic cortex; DP, dorsal peduncular cortex; CPu, caudate putamen; AcbC, accumbens nucleus, core; Acbsh, accumbens nucleus, shell; LSI, lateral septal nucleus, intermediate part; LSv, lateral septal nucleus, ventral part; LSD, lateral septal nucleus, dorsal part; VDB, nucleus of the vertical limb of the diagonal band; HDB, nucleus of the horizontal limb of the diagonal band; Acb, accumbens nucleus; MPA, medial preoptic area; MPO, medial preoptic nucleus; MnPO, median preoptic nucleus; BNST, bed nucleus of the stria terminalis; VLPO, ventrolateral preoptic nucleus; SFO, subfornical organ; LPO, lateral preoptic area; df, dorsal fornix; VP, ventral pallidum; PVT, paraventricular thalamic nucleus; Rt, reticular thalamic nucleus; LH, lateral hypothelamus; AHA, anterierhypothelamic area, anterior part; SO, supraoptic nucleus; SCN, suprechiasmatic nucleus; SLEA, sublenticular extended amygdala; ZI, zona incerta; RCH, retrochiasmatic area; RE, reuniens thalamic nucleus; Pe, periventriculorhypothalemic nucleus; CeA, central amygdaloid nucleus; MeA, medial amygdaloid nucleus; AHP, anterior hypothalamic area, posterior part; MePV, medial amygdaloid nucleus, posteroventral part; DMH, dorsomedial hypothalamic nucleus, VMH, ventromedial hypothalamic nucleus; CA1, field CA1 of hippocampus; CA3, field CA3 of hippocampus; PH, postiriorhypothelamic area; VLPAG, ventrolateral periaqueductal gray; LPAG, lateral periaqueductal gray; DMPAG, dorsomedial periaqueductal gray; DpMe, deep mesencephalic nucleus; RMC, red nucleus, magnocellular part; SNR, substantia nigra, reticular part; DR, dorsal raphe nucleus; PPTg, pedunculopontine tegmental nucleus; MnR, median raphe nucleus; PB, parabrachial nucleus; Mo5, motor trigeminal nucleus; RMg, raphe magnus nucleus; PCRtA, parvicellular reticular nucleus, alpha part; PnC, pontine reticular nucleus, caudal part; RVL, rostroventrolateral reticular nucleus; Sol, nucleus of the solitary tract. Scale bar: 500 μm.

**Figure supplement 3.**
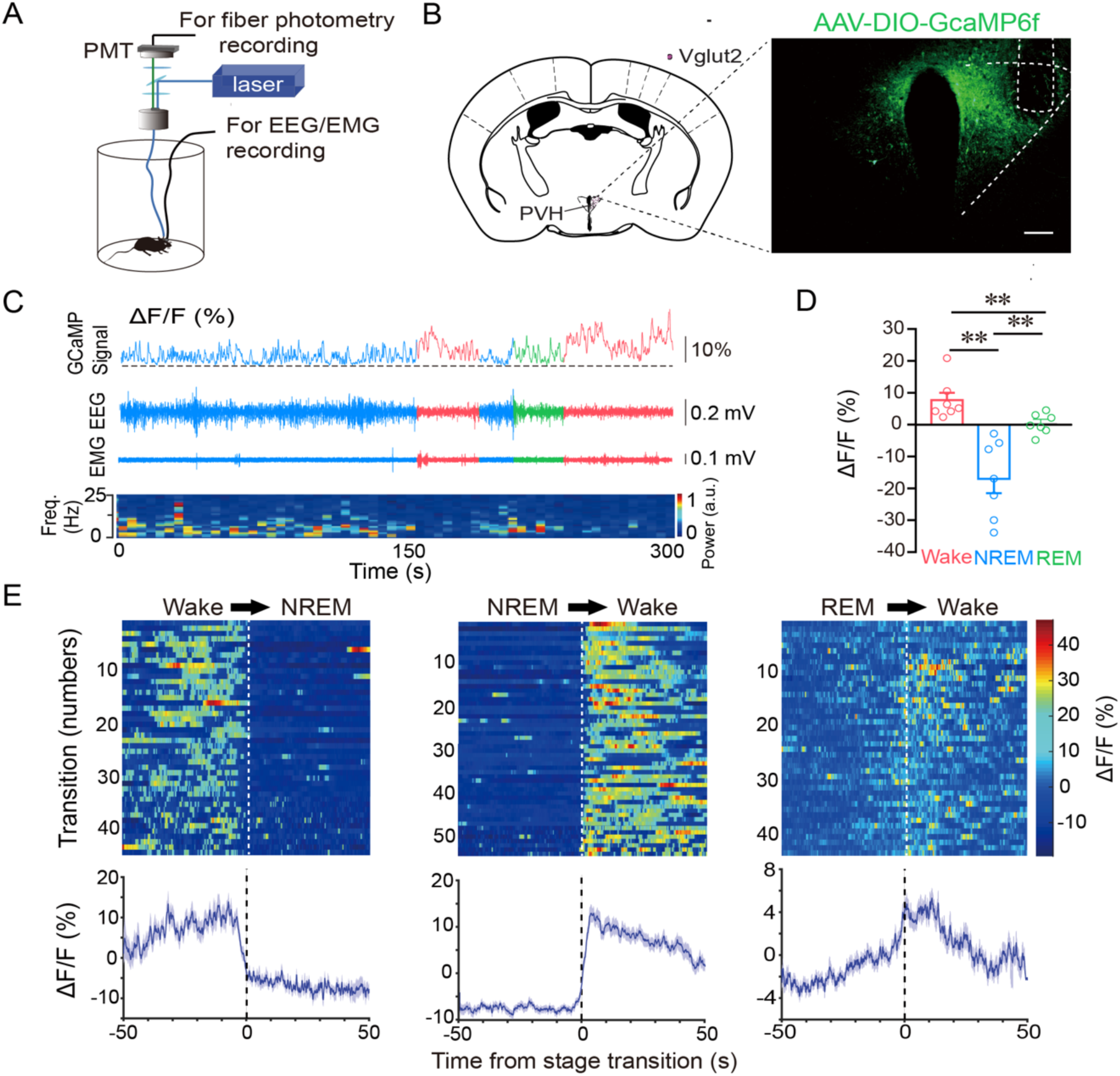
Population Ca2+ activity of PVH^vglut2^ neurons is coupled with wakefulness. **(A)** Schematic of the fiber photometry setup and *in-vivo* recording configuration (DM dichroic mirror, PMT photomultiplier tube). **(B)** Unilateral viral targeting of AAV-EF1α-DIO-GCaMP6f into the PVH, in which the tip of the fiber optic is above the PVH. Scale bar: 200 μm. **(C)** Representative fluorescent traces, relative EEG power, and EEG/EMG traces across spontaneous sleep–wake states. ΔF/F represents the change in fluorescence from the median of the entire time series. **(D)** Fluorescence (mean ± SEM) during wakefulness, NREM sleep, and REM sleep from three mice; the fluorescent signal was the highest during wakefulness, intermediate during REM, and the lowest during NREM sleep (n = 7 from 3 mice, one-way ANOVA followed by Tukey’s post-hoc tests; F_6,12_ = 2.94, *P*< 0.001; *P* [wake vs NREM] < 0.001, *P* [wake vs REM] < 0.001, *P* [NREM vs REM] = 0.013). **e** Fluorescent signals aligned to sleep–wake transitions. Upper panel, Individual transitions with color-coded fluorescent intensities (NREM to wake, n = 54; wake to NREM, n = 45; REM to wake, n = 44).

**Figure supplement 4.**
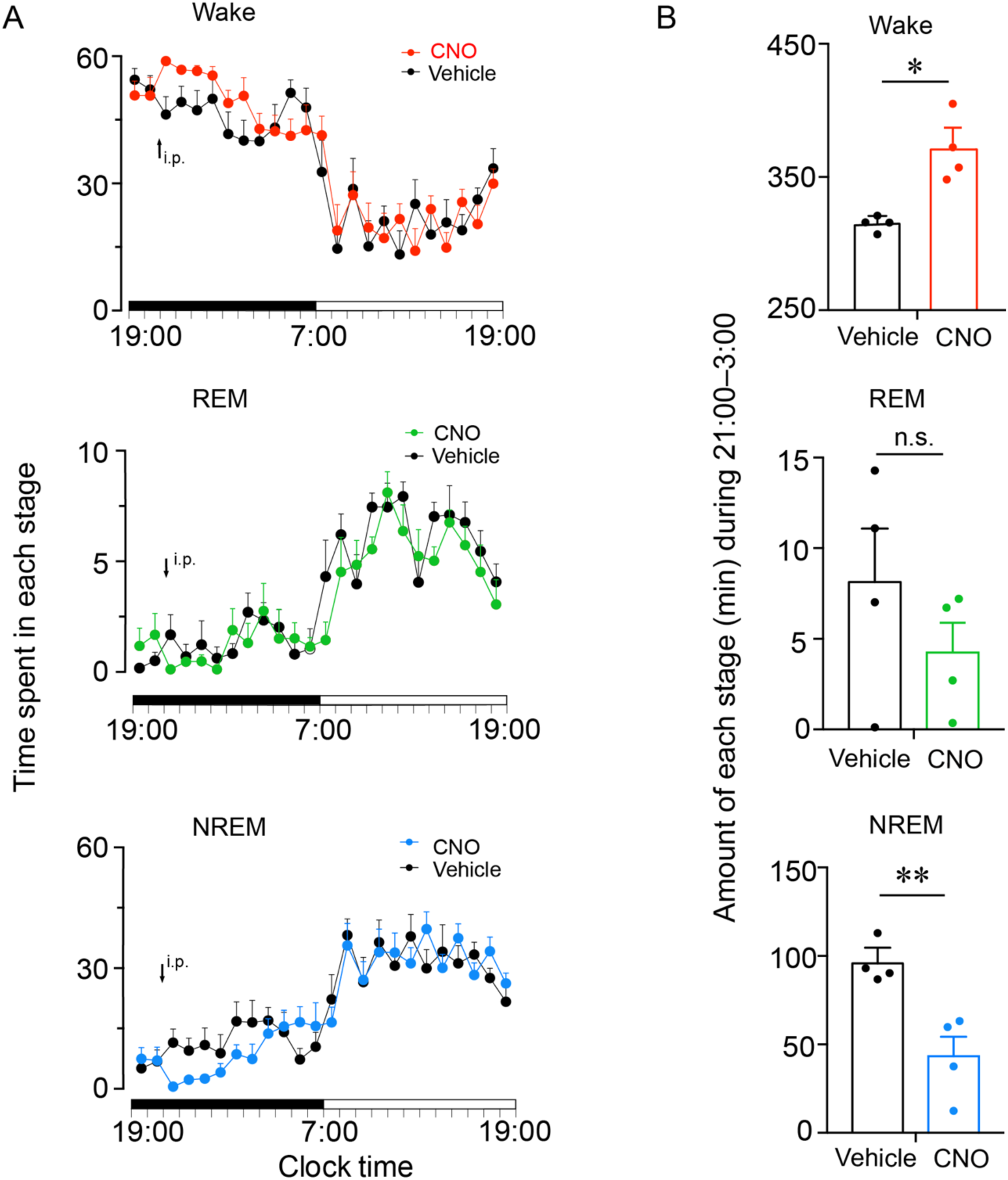
Chemogenetic activation of PVH^vglut2^ neurons during the dark phase increases wakefulness. **(A)** Time course of wakefulness, NREM sleep, and REM sleep following injection of vehicle or CNO in mice expressing hM3Dq receptor in PVH^vglut2^ neurons (n = 4). Statistical significance was determined using repeated-measures ANOVA. *P*>0.05. **(B)** Total time spent in each stage for 12 h after vehicle or CNO injection (n = 4). Statistical significance was determined using paired t test. \**P*< 0.05, \*\**P*< 0.01. n.s. denotes not significant.

**Figure supplement 5.**
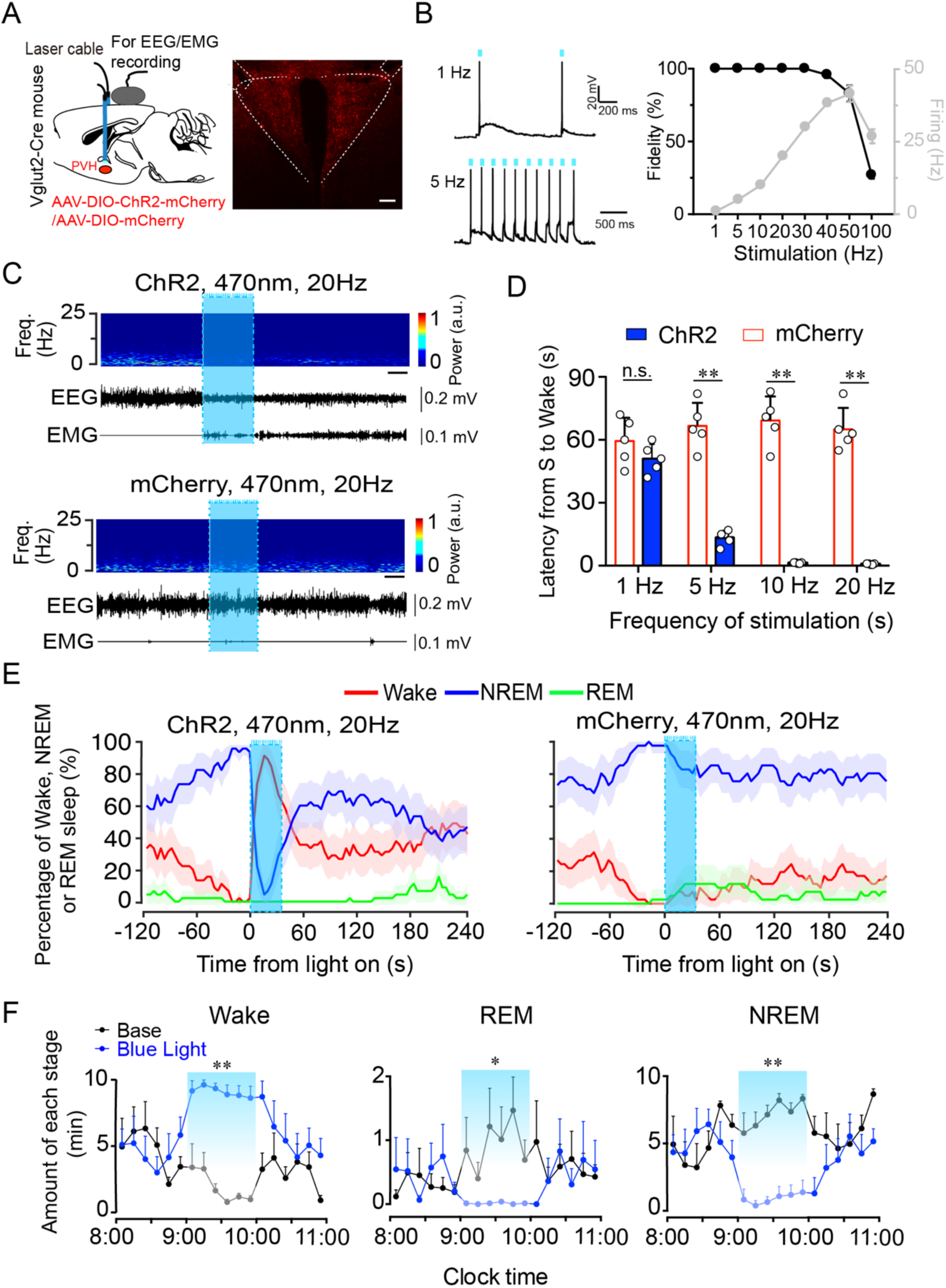
Optogenetic activation of PVH^vglut2^ neurons induces a rapid transition from NREM sleep to wakefulness. **(A)** Left: Schematic of optogenetic manipulation of PVH^vglut2^ neurons and EEG/EMG recordings. Right: ChR2-mCherry expression and location of optical fiber in the PVH. Scale bar: 200 μm. **(B)** Example traces (left) and fidelity of action potential firing (right) of ChR2-expressing PVH neurons evoked by 473-nm light stimulation with different frequencies. **(C)** Representative EEG/EMG traces, and heatmap of EEG power spectra showing that acute photostimulation (20 Hz/10 ms) applied during NREM sleep induced a transition to wakefulness in a ChR2-mCherry mouse. Scale bar: 10 s. **(D)** Latencies of transitions from NREM sleep to wakefulness after photostimulation at different frequencies (n = 5, unpaired t test; 1 Hz, t_8_ =1.4, *P* = 0.19; 5 Hz, t_8_ = 10.29, *P* < 0.01; 10 Hz, t_8_ =13.3, *P*< 0.01; 20 Hz, t_8_ =14.04, *P*< 0.01). **(E)** Sleep stage after blue-light stimulation in a PVH-vglut2-ChR2 mouse or PVH-vglut2-mCherry mouse. Percentages of NREM, REM, and wakefulness during short-stimulation experiments. **(F)** Time course during semi-chronic optogenetic experiments (20 Hz/10 ms, 25-s on /35-s off). The blue column indicates the photostimulation period of the stimulation group (n = 5, repeated-measures ANOVA; F_1,8_ = 59.37 (wake), 18.20 (REM), 103.30 (NREM); *P*< 0.001 [wake], *P* =0.003 [REM], *P*<0.001 [NREM]). Data represent the mean ± SEM (\**P* < 0.05, \*\**P*< 0.01).

**Figure supplement 6.**
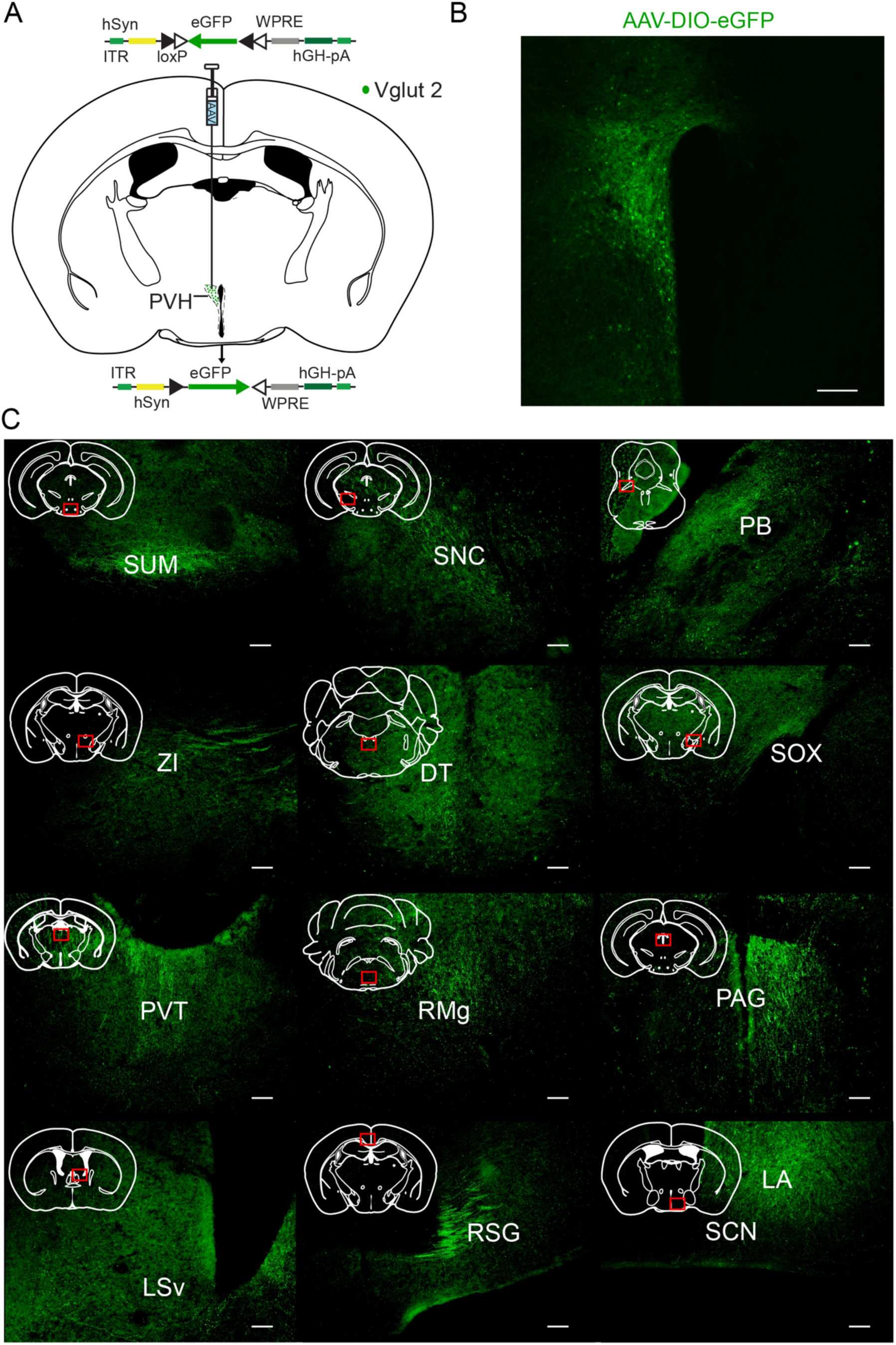
Representative regions with axonal projecting from PVH^vglut2^ neurons. **(A)** Schematic of the viral vectors and injection site for AAV-hSyn-DIO-eGFP in Vglut2-IRES-Cre mice. **(B)** Fluorescence images showing that AAV infection area in the PVH of a Vglut2-Cre mice. Scale bars: 200 μm. **(C)** Supramammillary nucleus, SUM; compact parts of substantial nigria, SNC; parabrachial nucleus, PB; zone incerta, ZI; dorsal terminal nucleus, DT. Supraoptic decussation, sox; paraventricular thalamic nucleus, PVT; raphe magnus nucleus, RMg; periaqueductal gray, PAG; ventral lateral septal nucleus, LSv; retrosplenielgranuler cortex, RSG; suprachiasmatic nucleus, SCN; lateralanterior hypothalamic nucleus, LA.

**Figure supplement 7.**
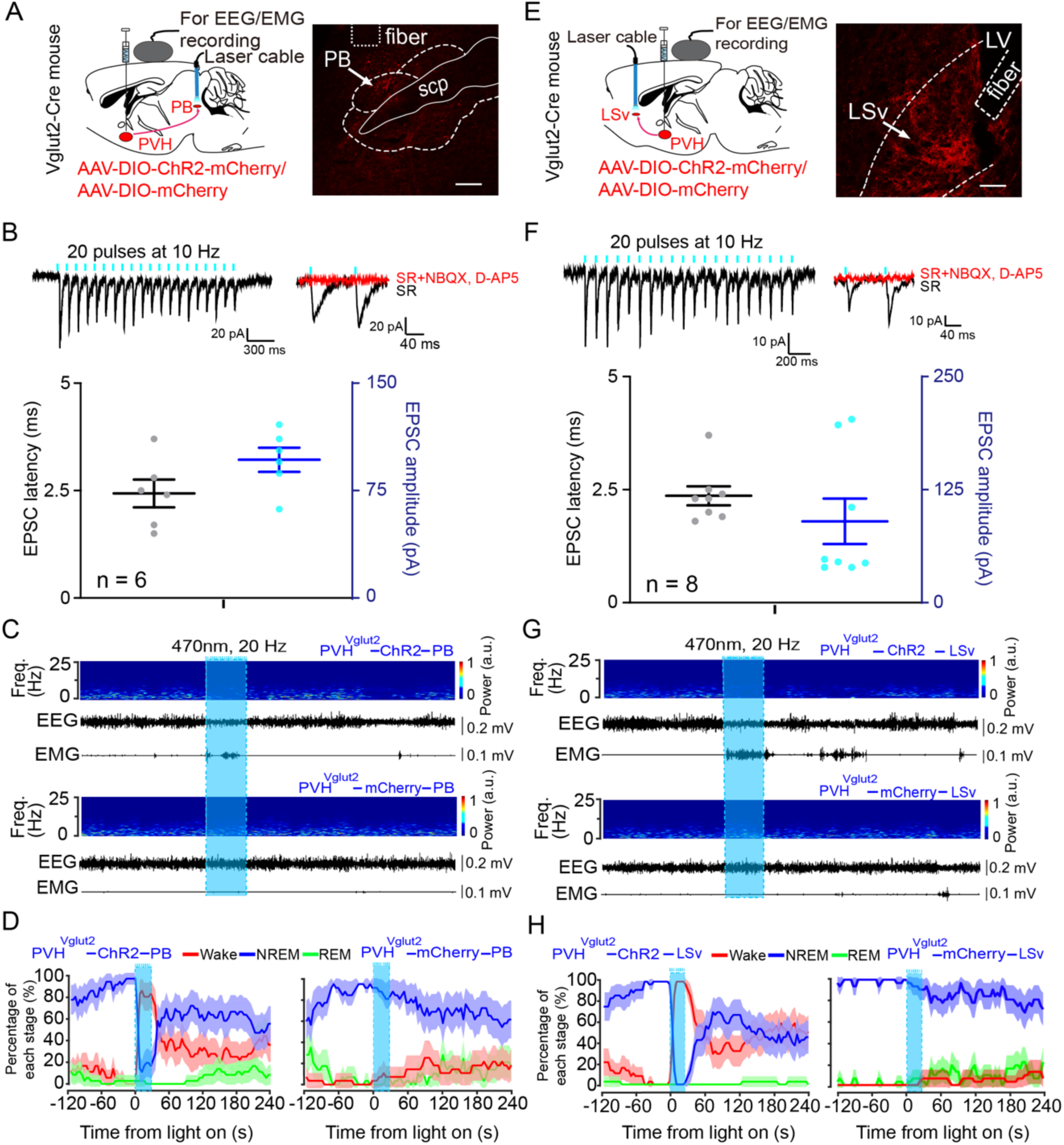
PVH^vglut2^ neurons control arousal through PB and LSv pathways. **(A, E)** Left: Schematic diagram showing the location of the optic fiber in the PB and LSv, and EEG/EMG recordings of a Vglut2-Cre mouse injected with AAV-ChR2-mCherry or AAV-mCherry in the PVH. Right: This brain section was stained against mCherry to confirm that ChR2 protein was expressed in the PVH. Scale bar: 200 μm. **(B, B)** Upper-left panel: Photostimulation-evoked EPSCs in PB neurons **(B)** and LSv neurons (**F**). Upper-right panel: Photostimulation-evoked EPSCs were completely blocked in the presence of NBQX (20 μM) and D-AP5 (25 μM). Lower panel: Latency (left axis) and amplitude (right axis) of light-evoked EPSCs in PB neurons (**B**) and LSv neurons (**B**). **(C, G)** Representative EEG/EMG traces, and a heatmap of EEG power spectra showing that acute photostimulation (20 Hz/10 ms) applied during NREM sleep induced a transition to wakefulness in a ChR2-mCherry mouse. Scale bar: 10 s. **(D, H)** Sleep stages after PB **(D)** and LSv **(H)** blue-light stimulation in ChR2-mCherry mice or mCherry control mice.

**Table supplement 1.**
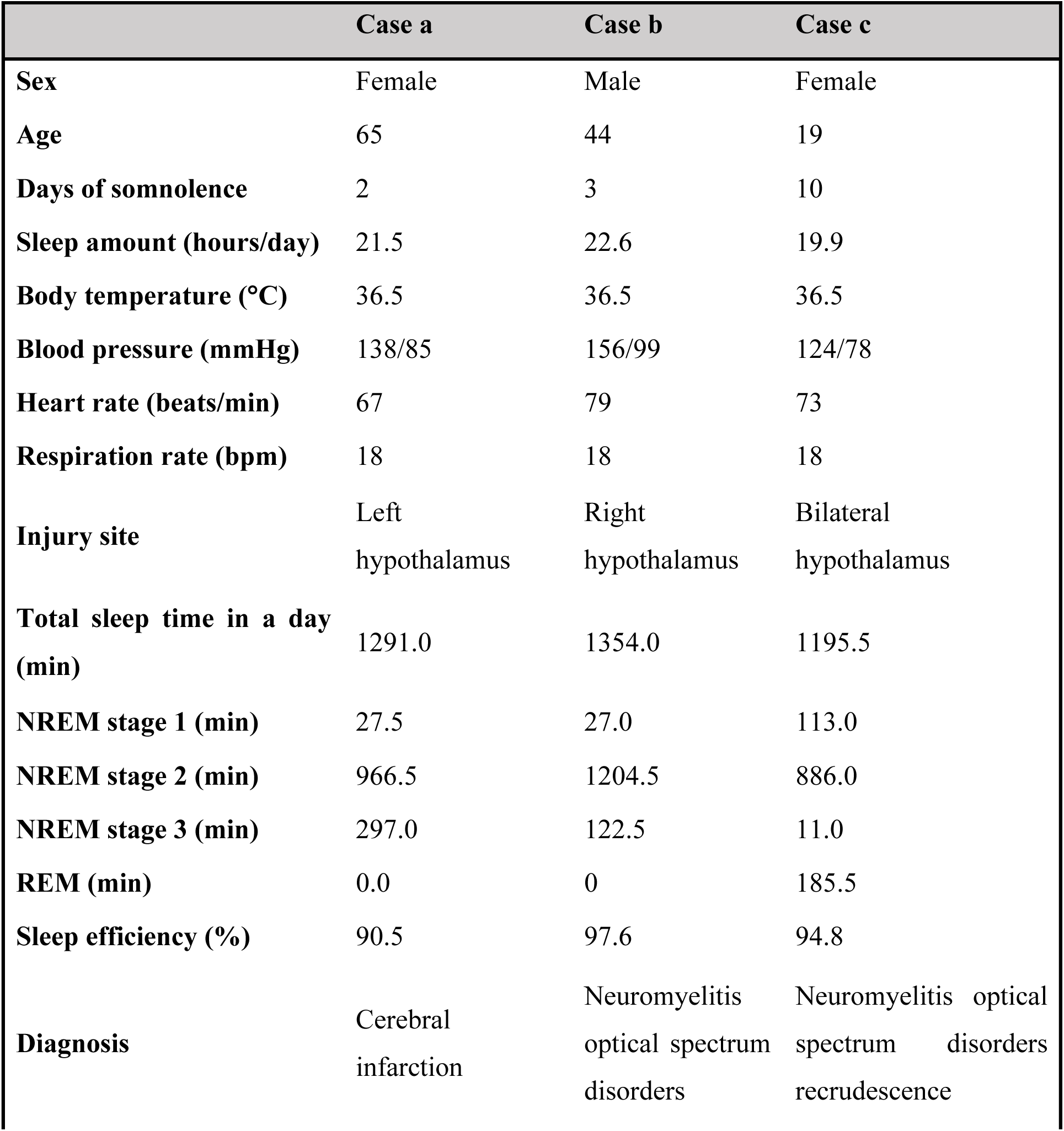
Clinical information of the included patients. Sleep efficiency (SE) is defined as “percentage of time slept of the total time spent in bed”. SE= total sleep time/total recording time * 100%.

|  | Case a | Case b | Case c |
| --- | --- | --- | --- |
| **Sex** | Female | Male | Female |
| **Age** | 65 | 44 | 19 |
| **Days of somnolence** | 2 | 3 | 10 |
| **Sleep amount (hours/day)** | 21.5 | 22.6 | 19.9 |
| **Body temperature (°C)** | 36.5 | 36.5 | 36.5 |
| **Blood pressure (mmHg)** | 138/85 | 156/99 | 124/78 |
| **Heart rate (beats/min)** | 67 | 79 | 73 |
| **Respiration rate (bpm)** | 18 | 18 | 18 |
| **Injury site** | Left<br>hypothalamus | Right<br>hypothalamus | Bilateral<br>hypothalamus |
| **Total sleep time in a day (min)** | 1291.0 | 1354.0 | 1195.5 |
| **NREM stage 1 (min)** | 27.5 | 27.0 | 113.0 |
| **NREM stage 2 (min)** | 966.5 | 1204.5 | 886.0 |
| **NREM stage 3 (min)** | 297.0 | 122.5 | 11.0 |
| **REM (min)** | 0.0 | 0 | 185.5 |
| **Sleep efficiency (%)** | 90.5 | 97.6 | 94.8 |
| **Diagnosis** | Cerebral<br>infarction | Neuromyelitis<br>optical spectrum<br>disorders | Neuromyelitis optical<br>spectrum disorders<br>recrudescence |

